# Emergence of Value and Action Codes in Bimodular Spiking Actor–Critic Networks

**DOI:** 10.1101/2025.11.26.690655

**Authors:** Carlos Antolín, Joan Falcó-Roget, Luis Serrano-Fernández, Néstor Parga

**Affiliations:** Departamento de Física Teórica, Universidad Autónoma de Madrid, 28049 Madrid, Spain; Centro de Investigación Avanzada en Física Fundamental, Universidad Autónoma de Madrid, 28049 Madrid, Spain; Computational Neuroscience Group, Sano Centre for Computational Medicine, Czarnowiejska 36, 30-054, Kraków, Poland

**Keywords:** Decision-making, value coding, economic choice, actor–critic architecture, reinforcement learning, spiking recurrent neural networks

## Abstract

Decision-making depends on coordinated computations distributed across dorsal and ventral circuits, often described as actor–critic systems. We examined how this division of labor gives rise to value- and decision-related representations by training a bimodular recurrent network on an economic choice task and transferring it to an expanded gated spiking model that preserved its latent dynamics while stabilizing the underlying neural representations. The two modules assumed complementary roles: the actor encoded action-value differences and spatial contingencies required for selecting between actions, whereas the critic represented object values predictive of reward. Neurons across modules were selective for object, action, total, and difference values, supporting decision- and confidence-related activity. Low-dimensional analyses revealed structured trajectories reflecting temporal evolution, spatial configuration, and value-dependent divergence. These results clarify how distributed circuits can jointly implement valuation and action selection, providing a foundation for linking reinforcement-trained recurrent models with anatomically inspired dorsal–ventral frameworks.

## Introduction

A central objective in systems neuroscience is to understand how distributed cortical and subcortical circuits support value-based decision-making [1, 2]. Among these circuits, fronto–striatal loops are often described as comprising a dorsal pathway associated with action selection and motor planning, and a ventral pathway associated with value representation and outcome evaluation [3]. This anatomical and functional division has motivated computational accounts based on reinforcement learning (RL) theory [4] in which dorsal structures operate as an actor, selecting actions according to learned policies, whereas ventral structures act as a critic, predicting expected reward and guiding learning [5–9].

Recurrent neural networks (RNNs)—typically trained with supervised learning—have been widely used to model perceptual decision-making [10–16]. Reinforcement learning methods have also been applied to train single RNNs for value-based behavior, showing that rich value and choice signals can emerge directly from recurrent dynamics [17–21]. Yet only a few studies have examined value-based decision-making in architectures that explicitly instantiate the dorsal–ventral division [17], and none has characterized the computations that emerge in such modular systems. As a result, the computational capabilities of minimally trained dorsal–ventral architectures, and their potential to generate value- and choice-related signals resembling those observed in fronto–striatal circuits [22–24], remain poorly understood. Understanding these capabilities is important because interacting actor–critic–like modules could, in principle, reveal how valuation, choice, and confidence-related signals arise from distributed recurrent dynamics. Our goal here is to characterize how a minimal dorsal–ventral architecture performs valuation and action selection when trained via reinforcement learning.

To investigate this, we trained a bimodular actor–critic spiking RNN on the economic choice task of Padoa-Schioppa and Assad [22]. This task, in which monkeys choose between two juice offers differing in quantity and identity, maps naturally onto the hypothesized functions of the critic (valuation of juice objects) and the actor (selection between spatial actions based on offer location). By analyzing how the trained network solves this specific task, we expect to gain direct insight into the computational capabilities and limitations of the bimodular architecture, and thereby develop intuitions about how it could be modified or extended to yield better descriptions of the processes involved in value-based decision-making.

More specifically, we adopted the actor-critic architecture defined in [17], trained with the REINFORCE algorithm [25]. Additionally, to probe the resulting dynamics at a more biologically realistic level, we developed a method to transfer a trained continuous RNN into an expanded gated spiking network. This mapping preserves the low-dimensional latent geometry of the original actor–critic system while enabling single-neuron and population analyses within a more biologically grounded spiking regime.

This framework allows us to pose several central questions about dorsal–ventral interactions during value-based choice. First, what neural subpopulations emerge in each module, and do they correspond to the object-value and action-value representations reported experimentally? Second, what decision- and confidence-related variables can be decoded from the activity of each module? Third, how does the actor resolve choices when it is more natural to make decisions in object-value space than in action-value space? Fourth, how do intermodular connectivity and directionality shape spatial, value-related, and decision-related signals? Finally, how are these computations expressed in low-dimensional population dynamics?

Our results show that the actor and critic modules develop distinct yet complementary coding schemes aligned with their reinforcement-learning objectives. The critic comes to represent object values predictive of reward, whereas the actor encodes action-value differences and spatial contingencies that support commitment to a particular choice. Across modules, we identify neural populations selective for object value, action value, total value, value difference, decision, and confidence. Furthermore, both networks exhibit low-dimensional, structured trajectories that branch according to spatial configuration and diverge along decision- and value-difference axes. These findings demonstrate that a minimal dorsal–ventral architecture can reproduce several hallmark computational motifs observed across fronto–striatal circuits, while also highlighting the architectural features that constrain specific value-based computations.

## Results

While recurrent neural networks (RNNs) have been successfully trained to perform perceptual decisionmaking tasks [10, 11, 13, 14], applications of trained RNNs to investigate value-based behavior remain comparatively rare [17–21]. Moreover, the internal structure and representational content of such models are often underexplored. Here we use a bimodular actor–critic architecture to study value-based decision-making in a framework inspired by dorsal and ventral fronto–striatal circuits.

We begin by describing the economic choice task [22] underlying our study. We then outline our method for transferring trained GRU networks into spiking recurrent networks of LIF-type neurons. Finally, we analyze the resulting spiking networks to characterize how subjective values, action values, choices, confidence signals, and key dynamical motifs emerge from the trained bimodular system.

### Task and behavioral structure

We focused on a classic economic choice task in which monkeys choose between two juice offers [22]. Each trial begins with a fixation period, followed by the simultaneous presentation of two offers representing varying amounts of juice a and juice b (Fig. 1a). Critically, juice identity and spatial position are dissociated; on each trial, the two juices are randomly assigned to left or right positions. After a variable delay, the agent selects one of three possible actions: maintain fixation, choose left, or choose right. The choice behavior is typically well captured by a sigmoid psychometric function relating choice probability to offer type (Fig. 1b), from which an indifference point *m*^*∗*^ (defined as the ratio at which the agent chooses either option with equal probability) can be estimated.

**Figure 1.**
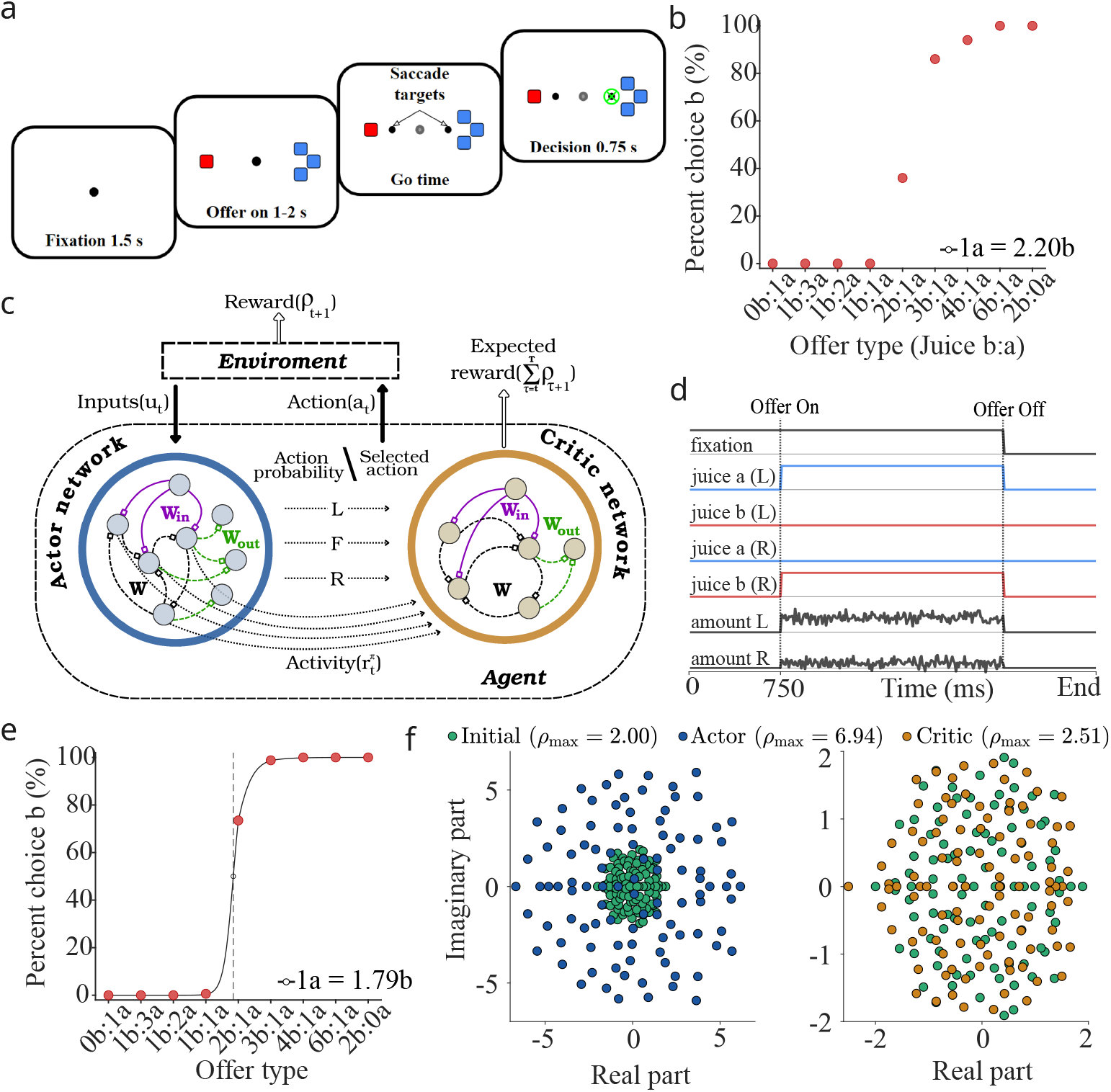
A gated recurrent actor–critic system for a value-based economic choice task. (**a**) Temporal structure of the task. Each trial began with a fixation cue, followed by the presentation of two juices (a and b) in variable amounts indicated by color and number of squares (e.g., 1 unit of juice b, red; 3 units of juice a, blue). The offer was shown for 1–2 s, after which the animal indicated its choice with a saccade and received the chosen juice. (**b**) Fraction of trials in which juice b was chosen for an example animal in [22]. The horizontal axis shows the offered ratios of juice a:b; the indifference point was 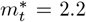. (**c**) Actor–critic architecture trained to perform the task. At each time step *t*, the actor (blue) received input *u*_*t*_ through non-plastic input weights (**W**_in_, solid purple) and selected one of three actions *a*(*t*) ∈{*fixate, left, right*}. The critic (brown) received the selected action and actor activity **r**^*π*^(*t*) via its own **W**_in_ to predict future reward *v*_**F**_. During fixation, the actor learned to choose *fixate* to avoid trial abortion (negative reward *ρ* = −1). During the decision period, rewards *ρ*(*t*) depended on the offered and chosen juices, after which the environment transitioned to a new state. Recurrent (**W**, dashed black) and output (**W**_out_, dashed green) weights were updated according to the respective learning rules (see Methods). (**d**) Inputs to the actor. A constant “fixation” input signaled the decision period onset. Juice identity and position were encoded by four inputs (blue/red for juices a/b), and juice amounts by two noisy inputs (gray). (**e**) Percentage of trials in which the trained system in **c** preferred juice b. The indifference point was *m*^*∗*^ = 1.79 (solid black; loss *L*_*A*_ = −0.3274). (**f**) Eigenvalue spectra of the recurrent weight matrices in the actor (left) and critic (right). Green: before training; blue (actor) and brown (critic): after training. Panels (**a**) and (**b**) adapted from [22]. Panel (**c**) adapted from [17].

### The GRU network system

To study this task, we build on the RNN architecture proposed in [17], which employs gated recurrent units (GRUs; [26]) arranged in an actor–critic configuration. In that framework, one GRU network (policy network) selects actions, and a second GRU network (value network) predicts expected reward based on the policy network’s internal state. In our simulations, we used a fixed set of nine offer types (Table 1), designed to span a range of value comparisons (see Methods for further details on the GRU system).

**Table 1.**
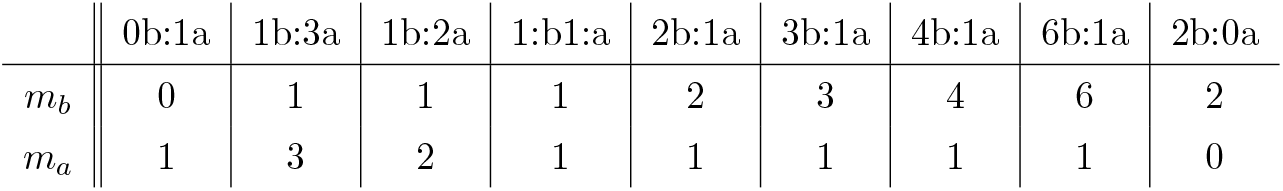
Offer types for the simulated task. Each column indicates an offer composed of *m*_*A*_ units of juice a and *m*_*B*_ units of juice b. The spatial arrangement of the offers was randomized across trials.

We first reproduced the bimodular GRU architecture of [17] (Fig. 1c), with intputs as described in Fig. 1d. Simulations were conducted using identical parameter values to those reported by the original authors (Methods). As a concrete case study, we selected a network trained on the economic choice task with a target indifference point 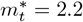, a condition examined both experimentally [22] (Fig. 1b) and in [17]. Fitting a sigmoidal function to the percentage of choices for juice *b* yielded an indifference point of *m*^*∗*^ = 1.79 for the GRU actor (Fig. 1e), to be compared with the target 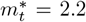 and the value *m*^*∗*^ = 2.0 reported in [17].

Training substantially modified the eigenvalue spectra of the recurrent weight matrices (Fig. 1f). The policy (actor) network showed a marked increase in spectral radius after training, suggesting richer dynamics that support flexible decision-making. In contrast, the value (critic) network exhibited a more moderate spectral expansion, remaining closer to the stability boundary, consistent with the requirement of stable dynamics for reliable reward prediction. These distinct dynamical regimes reflect the different computational roles of the two modules in the economic choice task.

### A bimodular system of gated spiking units

While the work in [17] used rate-based GRUs, our goal is to develop a more biologically plausible spiking counterpart that preserves the same latent dynamics. Our objective is to construct a system composed of two spiking recurrent neural networks (sRNNs; Fig. 2a) whose architecture and computational capacity mirror those of the bimodular GRU-based network. To this end, we adopt a gated leaky integrate-and-fire (gLIF) model that incorporates gating mechanisms analogous to those in GRUs. The following equations apply equally to both the actor and critic gLIF networks in Fig. 2a (see Methods for further details)

**Figure 2.**
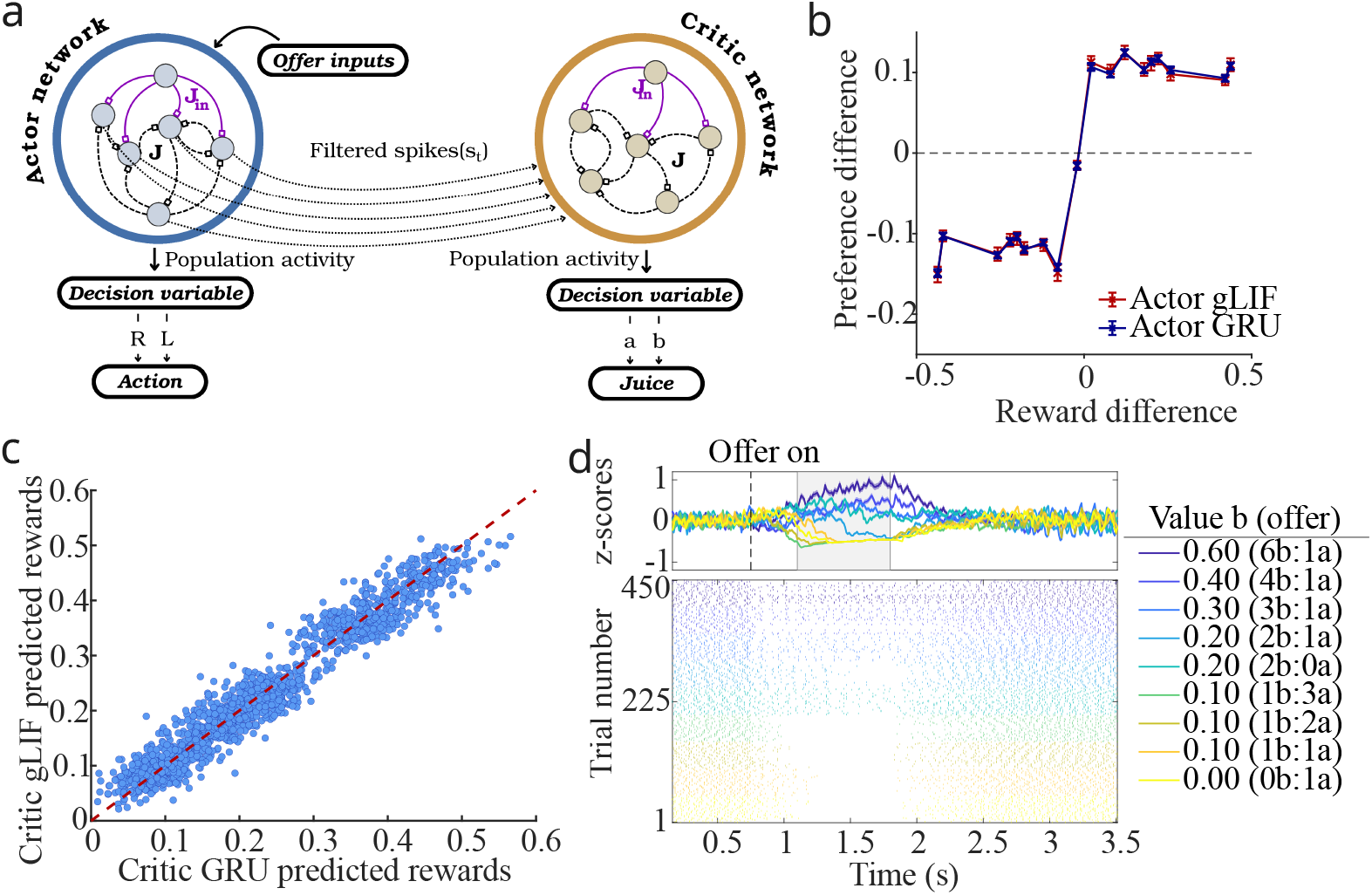
Bimodular system of gated spiking RNNs: scheme, optimization, and learning. **(a)** Schematic of the spiking actor–critic architecture after transferring the trained continuous RNNs. As in Fig. 1**c**, offers enter the actor network through random, untransferred input weights (**J**_in_, solid purple), and activity **s**(*t*) propagates through recurrent transferred weights (**J**, dashed black). The resulting activity is then passed to the critic via another set of random, untransferred weights, where it evolves through an analogous recurrent structure. In both modules, population activity defines a decision variable: left/right actions for the actor, and juice type a/b for the critic (see Methods). **(b)** Preferences of the trained actor GRU network (blue) and preferences decoded from the transferred actor gLIF network (red) as a function of reward differences between actions *R*_*R*_ − *R*_*L*_. Error bars denote the standard error of the mean across trials. **(c)** Trial-by-trial decoded rewards from the transferred critic gLIF network plotted against those from the critic GRU network. The red dashed line (*y* = *x*) indicates a perfect transfer. **(d)** Raster plot of a neuron from the transferred critic gLIF network tuned to the offered value of juice *b*, grouped by offer magnitude (color). The gray shaded area (1100–1800 ms) marks the analysis window used for sector classification. The Fano factor is 0.94, consistent with Poisson-like variability.

Spike trains generated by neuron *i* are first converted into continuous firing rates *s*_*i*_(*t*) through exponential filtering, modeling synaptic integration

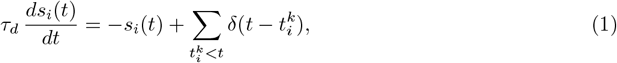

where *τ*_*d*_ = 100 ms is the synaptic decay time constant, and 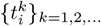 are the spike times. Collecting all units yields the vector of filtered spike trains **s**(*t*). The subthreshold membrane potential dynamics of each neuron follow

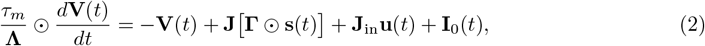

where **u**(*t*) is the external input vector, *τ*_*m*_ = 10 ms is the baseline membrane time constant, and **I**_0_(*t*) is a tonic input that sets the baseline firing regime (Methods). The vector **Λ** acts as a reset gate, modulating the effective time constants ***τ***_eff_ = *τ*_*m*_*/***Λ** and thereby controlling the rate of memory decay. The update gate **Γ** modulates the influence of recurrent input, acting as a multiplicative gain on the presynaptic firing rates **s**(*t*). These gates are defined as

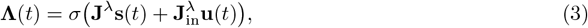

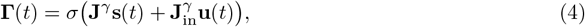

where *σ* is the element-wise logistic function, and **J, J**^*λ*^, **J**^*γ*^ and their corresponding input weights are learned transferred parameters. Although these gating variables do not correspond to specific anatomical mechanisms, gating-like computations have been proposed in prefrontal cortex [27] and provide a powerful means to dynamically regulate memory and information flow in cognitive models. We next describe how a trained bimodular GRU network is used to obtain a trained bimodular gLIF network.

### Learning transfer from a continuous RNN to a spiking RNN

Recent advances in training spiking recurrent neural networks have leveraged rate-based RNNs as auxiliary systems to guide learning. Strategies include using recurrent currents as targets [28], constraining spiking dynamics via a small number of latent factors extracted from the rate network [29], or directly using the recurrent weight matrix of the rate network [30]. In contrast to these rate-guided approaches, a different class of methods relies on surrogate gradients [31, 32], which optimize spiking recurrent networks by replacing the non-differentiable spike function with smooth approximations, thereby enabling gradient-based learning in sRNNs.

In our work, we adopt an approach based on the recurrent weight matrix but introduce key modifications to enhance robustness and flexibility. The method of [30] establishes a one-to-one correspondence between neurons in the rate and spiking networks, including output neurons responsible for signaling decisions. This strict mapping is constrained to networks of equal size and may not support robust learning transfer [29].

To overcome these limitations, we introduce two main innovations. First, we map each low-dimensional continuous GRU (with *M* units; Fig. 1c) into a higher-dimensional spiking network with *N > M* neurons (Fig. 2a). This expansion is performed while preserving the geometry of the latent space, i.e., maintaining the norms and angles between population activity vectors (see below). Second, rather than relying on a small subset of output neurons to assess performance, we analyze the gLIF networks for the presence of neural populations tuned to relevant latent features of the task and use these populations as readouts of decision and value variables.

Learning transfer is first applied to the actor gLIF network, and subsequently to the critic gLIF network. When transferring learning to the critic, it receives as input the activity of the already-optimized actor gLIF network (Fig. 2a). This is feasible because, once training is completed, information flows unidirectionally from actor to critic, with no feedback from the critic to the actor [17].

### Network size expansion

The original GRU actor and critic networks consist of *M* units with firing rates denoted by the *M*-dimensional vector **r**(*t*). Our goal is to transfer the learned dynamics from these rate-based networks (Fig. 1c) to spiking networks containing *N > M* units, whose firing rates are represented by the *N*-dimensional vector **s**(*t*) (Fig. 2a). We embed the low-dimensional activity of each GRU network into a higher-dimensional space using a linear transformation

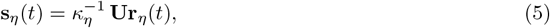

where *η* = *A, C* refer to the actor or critic, **U** is an *N × M* orthonormal matrix satisfying **U**^⊤^**U** = **I**_*M*_, and *κ*_*η*_ is a scaling factor that controls the overall firing rate level in the spiking network [30].

The recurrent connectivity in the spiking network is obtained by mapping the original *M × M* recurrent weight matrices **W, W**^*λ*^ and **W**^*γ*^ through the same transformation and applying the scaling factor *κ*_*η*_

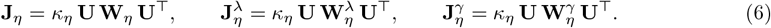

These matrices correspond, from left to right, to the recurrent weights of the activity network, the reset gate, and the update gate. Input weights are transformed as

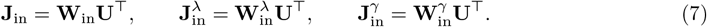

The transformation **U** preserves the geometric structure of the original dynamics (Supplementary Text S1.1). In particular, the relative norms and angles between activity vectors are maintained while increasing the network’s representational capacity. In a linear approximation [33], the temporal evolution and stability of the GRU and gLIF systems are governed by the same (although rescaled in the sRNNs) *M* nonzero eigenvalues *λ*_*i*_. The remaining *N* − *M* eigenvalues of the spiking connectivity matrix **J** are zero, corresponding to directions orthogonal to the image of **U**. Qualitatively, this expansion allows the latent variables driving the behavior of the GRU to be represented across a larger population of spiking units, providing a redundant, distributed code that enhances robustness to noise.

### Determination of scaling factors

The application of Eq. 6 is coupled to the determination of the scaling parameters *κ*_*A*_ and *κ*_*C*_. To transfer learning to the gLIF networks, we must identify the optimal values of these parameters, denoted by 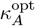 for the actor and 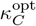 for the critic. This selection follows the same performance-based criteria used during GRU training: maximizing mean reward for the actor and minimizing reward prediction error for the critic. For the gLIF actor network, we minimize the loss

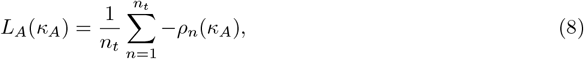

where *ρ*_*n*_(*κ*_*A*_) denotes the reward obtained on trial *n* as a function of *κ*_*A*_, and *n*_*t*_ is the number of trials used to estimate its mean. For the gLIF critic network, we minimize the squared prediction error

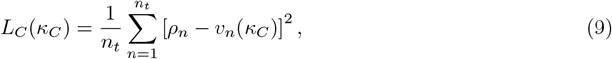

where *v*_*n*_(*κ*_*C*_) is the critic’s reward prediction for trial *n*, and *ρ*_*n*_ is the reward delivered following the actor’s choice.

### Learning transfer to the actor spiking RNN

Learning transfer from the actor GRU (Fig. 1c) to the actor gLIF network (Fig. 2a) proceeds as follows. For a given *κ*_*A*_, we estimate the loss in Eq. 8. To compute the reward *ρ*_*n*_(*κ*_*A*_) on each trial, we first infer the gLIF actor’s choice. This is done by constructing a linear readout of the preference difference between actions R and L. The readout is trained via regression on a dataset comprising (i) the preference-difference output of the actor GRU and (ii) the corresponding firing-rate vectors of the actor gLIF network in a fixed temporal window after offer onset (Fig. 2d; Methods). Once trained, the readout is used to infer the choices made by the actor gLIF network in *n*_*t*_ simulated trials, from which we compute the corresponding rewards and evaluate the loss as a function of *κ*_*A*_. By repeating this procedure across a range of candidate scaling factors, we identify an interval where the loss is minimized. A subsequent fine–tuning step (Methods) yields the optimal scaling value, 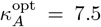. To assess the quality of the transfer, we compared the preference-related output of the actor GRU with the decoded preference from the actor gLIF network as a function of reward difference. The gLIF actor closely reproduced the preference–reward relationship observed in the GRU (Fig. 2b). A consistent result was obtained from the readout-based choice pattern, whose indifference point (*m*^*∗*^ = 1.84) closely matched the GRU estimate (*m*^*∗*^ = 1.79; Supplementary Fig. 1).

### Learning transfer to the critic spiking RNN

Once the actor gLIF network is obtained, it sends its spiking activity to the critic gLIF network (Fig. 2a). Learning transfer to the critic follows a procedure analogous to that used for the actor, but now using the loss in Eq. 9. For a given *κ*_*C*_, we train a linear readout that maps the critic gLIF activity to reward predictions. The readout is trained on pairs consisting of the GRU critic’s reward predictions and the corresponding gLIF critic firing-rate vectors in the same trials. After training, we simulate *n*_*t*_ trials, apply the readout to estimate *v*_*n*_(*κ*_*C*_) in each trial, and evaluate the loss. Minimizing this loss yields an optimal 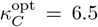 (Methods). Predicted rewards from the spiking critic closely match those of the GRU critic on a trial-by-trial basis, clustering along the identity line (Fig. 2c), indicating accurate transfer of the reward-prediction function. As an illustration of single-unit activity, Fig. 2d shows a critic neuron selective for the offer value of juice b, with Fano factor close to the Poisson regime.

After learning transfer, the spiking actor–critic system is capable of autonomously generating decisions for arbitrary offer pairs. One approach is to use the preference-difference readout from the actor gLIF network, which now operates independently of the GRU actor. A more principled and biologically interpretable approach, however, is to decode decisions directly from intrinsic population codes in the spiking actor and critic. We therefore next identify neural populations tuned to action values in the actor and to object values in the critic, and analyze how their activity supports decision-making.

### Action-value representations in the actor network

To study action-value coding in the actor gLIF network, we search for neurons tuned to the values associated with actions R and L. This requires care, because in the task design, juice identity and spatial position are dissociated [22]. Thus, decoding decisions in terms of action values is only meaningful when juice positions are explicitly taken into account on each trial.

For each neuron in the actor network, we perform a multilinear regression with the chosen action, the action values (defined as the reward magnitudes associated with the juices offered on the right or left [23]), and juice position as regressors (Methods). We then project each neuron’s regression coefficients for the R and L action values, *β*_2_ and *β*_3_, into a two-dimensional action-value space. The resulting vectors characterize the direction and magnitude of each neuron’s tuning relative to action values. Following [34], we partition this space into eight angular sectors of 45^*°*^ (Fig. 3a; Methods). Neurons with vectors aligned with the horizontal (vertical) axis predominantly reflect value for action R (L). Vectors near 135^*°*^ or 315^*°*^ indicate encoding of the signed difference between action values, while orientations around 45^*°*^ or 225^*°*^ are associated with net action value signals that sum across actions.

**Figure 3.**
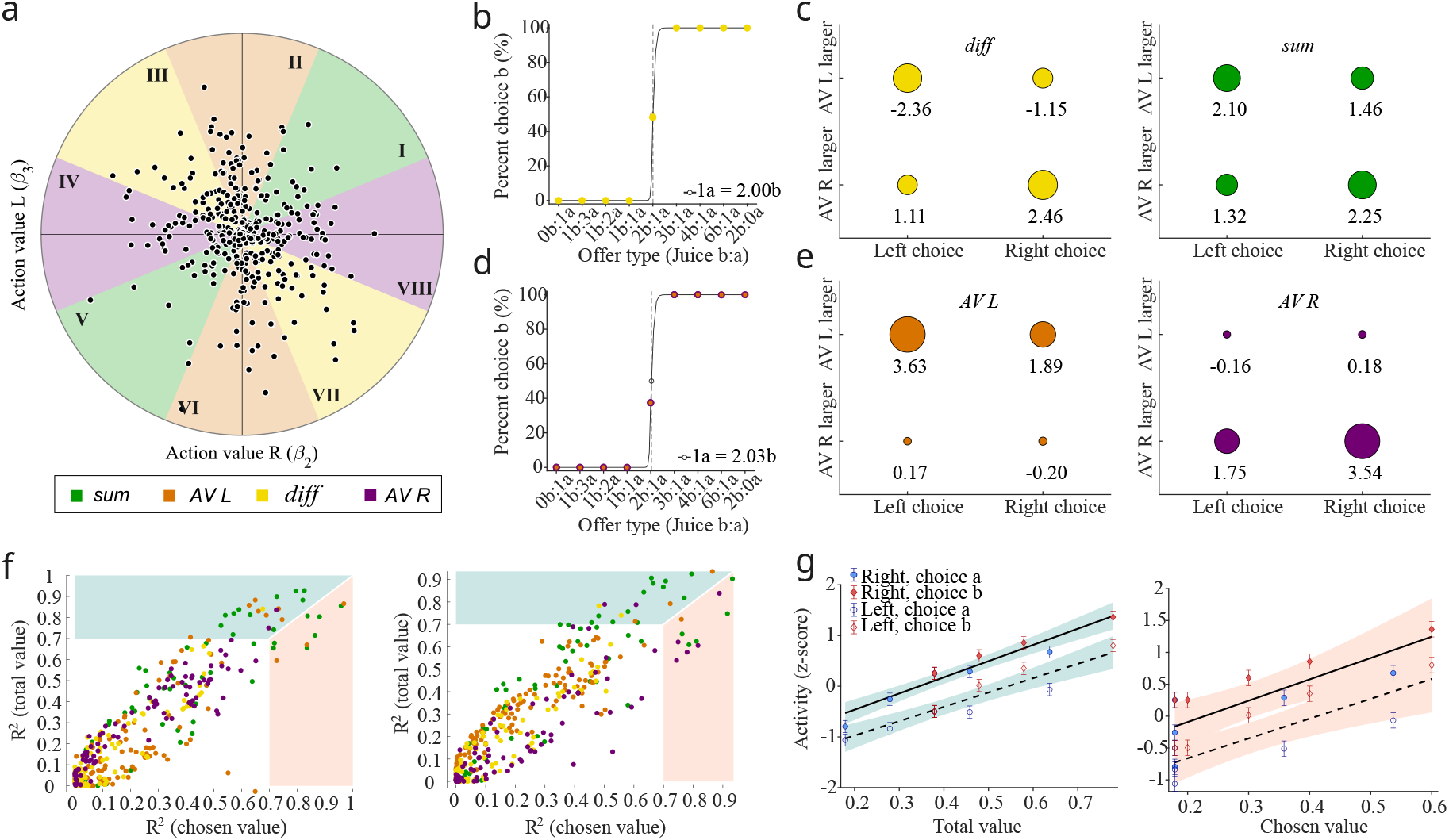
Neural populations and decision variables in the actor gLIF network. **(a)** Classification of actor neurons in action-value space. Neurons with significant action-related encoding (F-test, *p <* 0.01) were assigned to *sum* (*N* = 69), *diff* (*N* = 99), *AV R* (*N* = 97), or *AV L* (*N* = 125) populations according to the polar angle *θ* derived from regressors (*β*_2_, *β*_3_). **(b)** Choice pattern derived from the decision variable 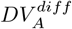 (difference neurons; sectors 3 and 7). Sigmoid fit: *m*^*∗*^ = 2.00,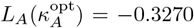. **(c)** Averaged partial residuals for *sum* and *diff* neurons. Trials were grouped by the action predicted by 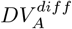 and whether the corresponding action value exceeded the alternative (see Methods). Circle radii scale with the absolute value of the residuals. **(d)** Choice pattern from the decision variable 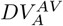 (action-value neurons; sectors 2, 4, 6, 8). Sigmoid fit: *m*^*∗*^ = 2.03, 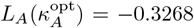. **(e)** Same as in **c** but using action-value neurons and the variable 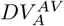. **(f)** Coefficients of determination (*R*^2^) for chosen vs. total value encoding for each neuron; colors indicate sectors as in **a.** Teal/coral shading marks neurons better explained by total/chosen value, respectively. **(g)** Linear regressions of total (teal) and chosen (coral) value against population-averaged activity in the *sum* population (sectors 1 and 5). Solid/dashed lines correspond to juice a on the right/left. Total value explains more variance 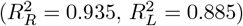 than chosen value 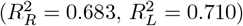. Shaded areas: 95% CIs. Error bars: mean *±* SEM.

Each action-value-related pattern corresponds to two mirror sectors with opposite signs. We refer to the corresponding neural populations as plus and minus populations (e.g., sectors III and VII are the *diff* + and *diff* – populations, respectively; Fig. 3a).

Neurons in the *diff* populations are natural candidates for mediating action selection [24]. Accordingly, we define a decision variable 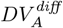 from compounded partial responses of these neurons (Methods). A positive value of 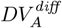 corresponds to a right choice, and a negative value to a left choice. The resulting choice pattern obtained using this decision variable (Fig. 3b) exhibits an indifference point at *m*^*∗*^ = 2.00, matching well its target value.

To test whether trials classified as R choices by this decision variable correspond to *AV* _*R*_ higher than AV_*L*_, and conversely for L choices, we examine how action values modulate responses in the *diff* populations. Following [23], we computed partial residuals, averaged across trials and neurons in the *diff* populations, grouped by the chosen action and whether *AV* _*R*_ exceeded *AV* _*L*_. Activity in this population reflects the network’s decision: stronger signals for action R are associated with R choices and vice versa, confirming that the population is differentially activated depending on the strength and direction of the preferred action (Fig. 3c).

Although the *diff* population is the most direct substrate for decoding the actor’s choice, populations tuned selectively to *AV* _*L*_ (sectors II and VI) or *AV* _*R*_ (sectors IV and VIII) can also be used. By taking the difference between their activities (Methods), we obtain an alternative decision variable 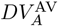, which produces a choice pattern with an indifference point *m*^*∗*^ = 2.03 (Fig. 3d). Analyzing the residual activity of these sectors shows that they exhibit stronger responses when the chosen action matches their preferred direction (Fig. 3e).

Neurons in the *sum* sectors of the actor (Fig. 3c, panel *sum*) do not distinguish individual action values. Their tuning could reflect the total action value (*AV* _*R*_ + *AV* _*L*_), or the chosen action value. Because these quantities are correlated, we fit, for each neuron, separate linear regressions using total or chosen action value as predictors, and compare coefficients of determination *R*^2^ [22] (Methods). For the configuration in which juice *a* is presented on the right, 13 neurons are selective only to total value and 3 only to chosen value, while 27 encode both; among the latter, total value explains activity better in most cases (19 vs. 8; Wilcoxon signed-rank test, *p* = 0.003). For the left configuration, 10 neurons are selective only to total value and 4 only to chosen value, with 26 encoding both; again, total value accounts for a larger fraction of variance (18 vs. 8; *p* = 0.022). At the population level, mean activity of *sum* neurons across trials is better captured by total action value 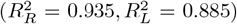 than by chosen action value 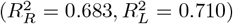 (Fig. 3f,g). These findings indicate a predominance of total-value signals in the *sum* population, consistent with the idea that the actor integrates action values into a global measure of the desirability of available actions, like in *sum* sectors in experimental settings [24, 34].

### Juice-value representations in the critic network

In the original bimodular GRU system, the actor was explicitly trained to make decisions, while the critic learned to provide a baseline value signal [17]. However, in the present task, decisions can naturally be framed in terms of object (juice) values, as extensively explored in neural recordings [22, 35]. Given the dissociation between juice identity and position in the task design, object-value-based decision-making is a natural computational strategy. We therefore asked whether neurons in the critic gLIF network also support decision-making in juice-value space.

We first identify neurons whose activity is significantly modulated by the values of the offered juices. For each neuron, we fit a multilinear regression model with the values of juices a and b as independent variables and include juice position as an additional regressor to test position dependence (Methods). Each neuron’s response is then represented as a vector in a two-dimensional object-value space, with components given by the regression coefficients for the two juices [34] (Fig. 4a).

**Figure 4.**
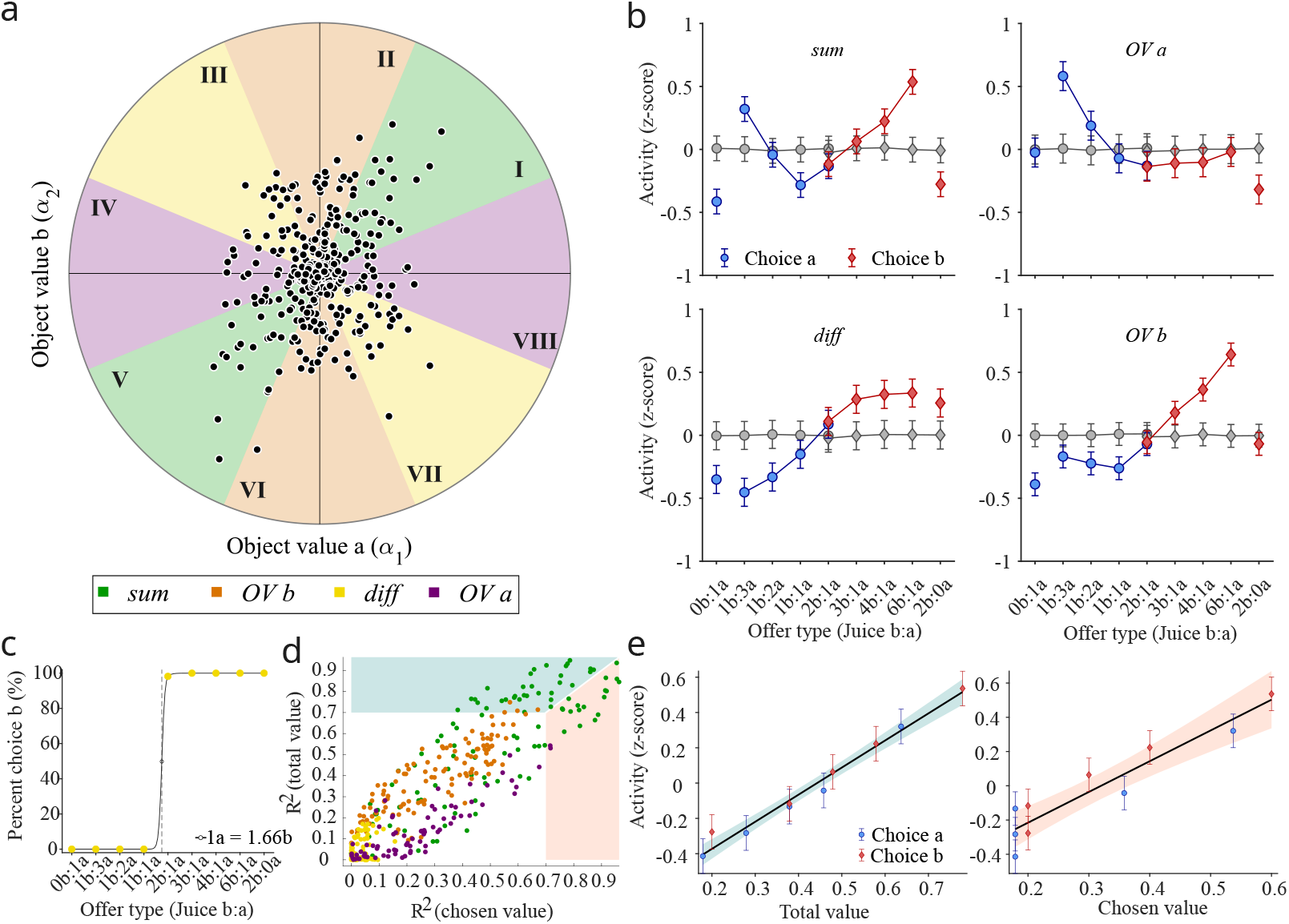
Neural populations and decision variables in the critic gLIF network. **(a)** Classification of critic neurons in object-value space using the polar angle *θ* from regressors (*α*_1_, *α*_2_). Populations: *sum* (*N* = 104), *diff* (*N* = 81), *OV a* (*N* = 76), and *OV b* (*N* = 122). **(b)** Mean tuning curves for each population. Activity was Z-normalized, sign-flipped when needed (sectors 3–6), and split by choice (blue/red for juice a/b). Gray points: pre-offer baseline. **(c)** Choice pattern from 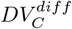 (difference neurons, sectors 3 and 7). Sigmoid fit: *m*^*∗*^ = 1.66,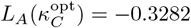. **(d)** Coefficients of determination (*R*^2^) for chosen vs. total value encoding. Shaded regions mark neurons better fit by total (teal) or chosen (coral) value. *Sum* neurons (green) show a slight bias toward total value. (**e**) Linear regressions of total (teal) and chosen (coral) value against activity in the *sum* population (sectors 1 and 5). Total value explains more variance (*R*^2^ = 0.978) than chosen value (*R*^2^ = 0.889). Shaded areas: 95% CIs; Error bars: mean *±* SEM.

As in the actor analysis, we divide this space into eight angular sectors of 45^*°*^, following previous approaches to value-related encoding [24, 34]. The angle of each neuron’s tuning vector provides a geometric descriptor of its computational role. Vectors aligned with the horizontal or vertical axes indicate selective sensitivity to the value of a single juice (object-value coding). Vectors near 45^*°*^ or 225^*°*^ reflect coding of the *sum* of juice values (total value), often associated with motivational drive. Vectors around 135^*°*^ or 315^*°*^ correspond to the signed difference in juice values, a canonical decision variable. Tuning directions near the unsigned value-difference axis may reflect confidence-related signals [24]. This geometric partition thus reveals structured functional specializations within the critic.

To test these interpretations, we examine average firing rates across offer types for each population (Fig. 4b). Neurons tuned to the value difference between juices (sectors III and VII) are natural candidates for signaling choice. We combine these neurons to define a decision variable 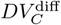, which we use to reconstruct the behavioral choice pattern (Methods). The resulting psychometric curve (Fig. 4c; *m*^*∗*^ = 1.66) is compatible with the actor-based choice patterns (Fig. 1e and Fig. 3d), indicating that the critic’s *diff* population also contains sufficient information to support decisions.

Neurons in the horizontal and vertical sectors exhibit tuning consistent with object-value encoding for juices *a* and *b*, respectively (Fig. 4b, panels OV a and OV b), similar to the object-value signals reported in OFC [22]. We then focus on the critic’s *sum* sectors (sectors I and V), which display a characteristic U-shaped response profile across offer types (Fig. 4b, panel *sum*), suggestive of total or chosen value coding. As with the actor, we compare total and chosen value models for each neuron using separate regressions and *R*^2^ (Methods). Most neurons with *R*^2^ *>* 0.7 for either predictor are found in the *sum* sectors (Fig. 4d, green dots). Among the 61 neurons sensitive to both variables, 35 are better explained by total value and 26 by chosen value (Wilcoxon signed-rank test, *p* = 0.043); an additional 11 neurons are selective only to total value and 8 only to chosen value. As expected, *diff* neurons show the weakest correlations with either predictor. At the population level, the mean activity of *sum* neurons across trials is also better captured by the total value (*R*^2^ = 0.98) than by the chosen value (*R*^2^ = 0.87; Fig. 4e). Thus, in the critic network, *sum* sectors predominantly encode total offer value, making total value the dominant signal in this population.

We have shown that both actor and critic networks can make decisions, yielding similar choice patterns (compare Fig. 3b and Fig. 4c). For offers with unambiguous decisions, the choices made by the two networks are highly consistent. For the split offer, where the two options have nearly identical values, each network selects either option with roughly equal probability, implying that their decisions should coincide only at the chance level. To test this, we computed the fraction of split trials in which the two networks selected the same option. This fraction was close to 50%, indicating that their choices were effectively uncorrelated in this ambiguous condition.

### Decision confidence

Confidence, commonly defined as the probability that a selected option is the better choice [36], has been linked to neural activity in several decision-related areas. In perceptual decision-making, confidence correlates with firing rates in lateral intraparietal cortex [37] in monkeys and OFC [38] in rodents, and can be formalized as the posterior probability of being correct in evidence-accumulation models [39]. In value-based choice, the dorsolateral prefrontal cortex encodes confidence-related quantities such as the unsigned value difference between options, while the signed value difference predicts choice [24].

Although confidence was not explicitly trained, we hypothesized, following [24], that the absolute value of the critic’s decision variable, 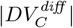, reflects an internal estimate of confidence. A simple argument from signal detection theory supports this interpretation [38]. If the values *V*_*a*_ and *V*_*b*_ of the two juices are corrupted by independent Gaussian noise with standard deviation *σ*, the probability of selecting the better option is

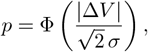

where |Δ*V* | = |*V*_*a*_ − *V*_*b*_| and Φ(·) is the cumulative distribution function of the standard normal distribution (Supplementary Text S1.2). We found that the absolute value of the normalized response of the *diff* population (absolute z-scores) follows the sigmoid-like relationship with |Δ*V* | predicted by *p*, indicating that the network encodes confidence on a trial-by-trial basis (Pearson *ρ* = 0.90, *p* = 0.001; Fig. 5a). When plotted as a function of offer type, this signal exhibits the characteristic V-shaped profile, with lowest confidence for the split offer (Fig. 5b, blue symbols).

**Figure 5.**
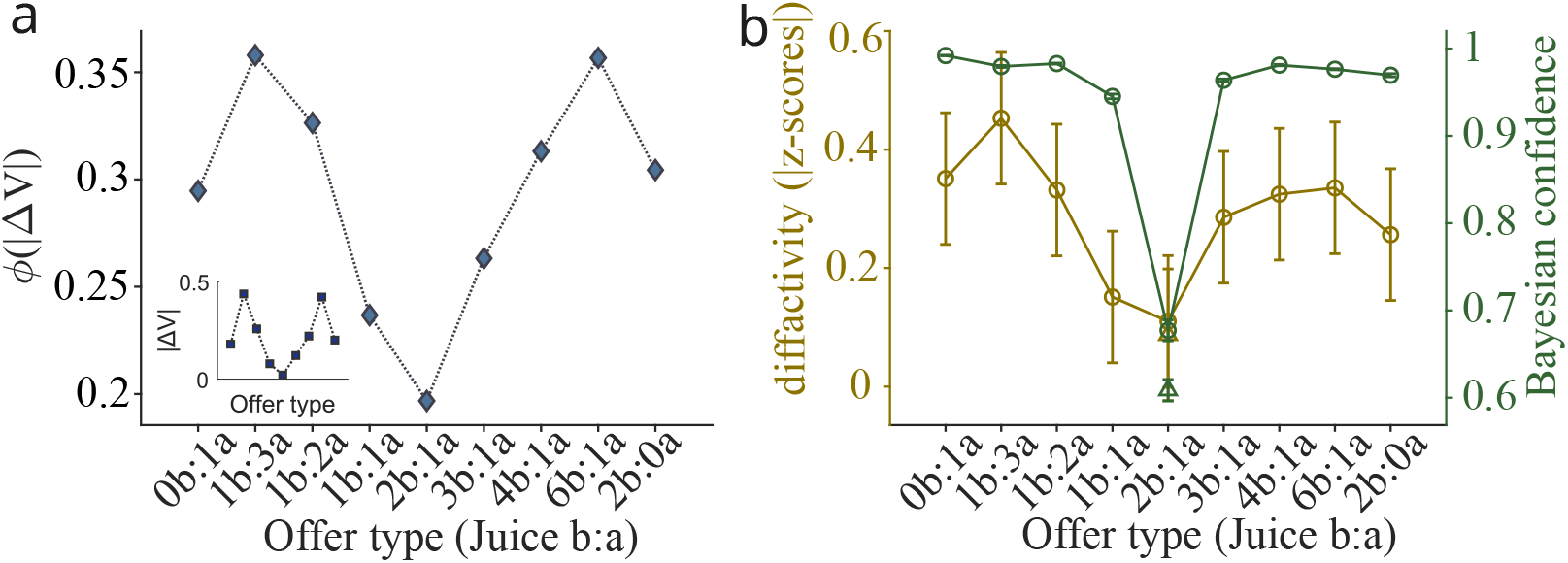
Neural activity in the *diff* population correlates with confidence. **(a)** Confidence function *ϕ*(|Δ*V*|) from signal detection theory, relating difficulty (absolute value difference |Δ*V*|) to confidence. Inset: distribution of |Δ*V*|. Activity of the *diff* population strongly correlates with *ϕ*(|Δ*V*|) (Pearson *ρ* = 0.90, *p* = 0.001). **(b)** *Diff* population activity (left axis) and Bayesian confidence (right axis), across offers. Circles: correct trials; triangles: errors. Activity closely followed with Bayesian confidence (Spearman *ρ* = 0.80, *p* = 0.013).

We also trained a dedicated readout neuron to estimate confidence from the *diff* population using Bayesian logistic regression [40] (Methods). The readout learned to predict the probability that the chosen option was correct, based on population activity. The resulting confidence estimates show the same V-shaped dependence on offer values (Fig. 5b, red symbols), and closely track those derived from the absolute activity of the *diff* population (Spearman *ρ* = 0.80, *p* = 0.013). Together, these results indicate that the critic network spontaneously develops a representation of decision confidence, even though confidence is not an explicit training target.

### Juice-position coding

Regression analyses in both networks revealed significant coding of juice position: 97% of actor neurons and 94.8% of critic neurons showed a significant position regressor. While position dependence is expected in the actor, it is more surprising in the critic, given the position-invariant value coding reported in OFC [22]. To examine this discrepancy, we analyzed how neural activity in each network depends on the spatial configuration of the offers.

We first asked whether neurons that show significant activity when juice *a* is presented on the right are the same as those active when the same juice is presented on the left. Within each of the two *diff* sectors of the critic, we computed the fraction of neurons that are significantly active in both spatial configurations. In the *diff* + sector (42 neurons), 40.5% were significantly active when juice *a* was on the right, 45.2% when it was on the left, and only 2.4% in both conditions. In the *diff* – sector (39 neurons), 35.9% were active for juice *a* on the right, 53.9% for juice *a* on the left, and none in both. This result indicates that decision-related activity in the critic occupies different regions of state space depending on spatial configuration, revealing a strong position dependence of *diff* representations.

We performed the same analysis in the actor. Here, the fraction of neurons encoding action value in both spatial configurations is noticeably higher. For example, the overlap reaches 15.1% in *diff* + and exceeds 26.3% in *AV* _*R*_+. This suggests that the actor develops partially position-invariant action-value codes required for mapping value onto spatially defined actions, whereas the critic refines the actor’s input to compute juice values, introducing additional spatial specificity.

Across the remaining sectors, a similar pattern emerged. In the critic, most neurons responded to only one spatial configuration, with only a small fraction tuned to both positions of juice *a* (Supplementary Table 1). In the actor, neurons selective for both configurations were slightly more frequent, but, as in the diff sectors, responses were still predominantly tied to a single spatial location for each juice (Supplementary Table 2).

### Dynamical organization of actor and critic networks

Both actor and critic possess sufficient neuronal and circuit-level resources to perform decisions, but they do so according to different computational objectives. The actor selects actions that maximize expected reward, whereas the critic predicts reward outcomes based on the actor’s internal state. Importantly, during training, juice identities and positions were randomized independently across trials, so the actor could not rely on fixed identity–action associations. Instead, it had to dynamically combine value and spatial information to generate the correct action on each trial.

To understand how these computations unfold dynamically, we analyzed population trajectories in neural state space using principal components analysis (Methods). In the actor, the geometry of trajectories provides direct evidence for how value and position are integrated (Fig. 6a). Before offer onset (squares), trajectories corresponding to different offers and positions evolve in an overlapping manner in state space. After offer onset, trajectories split into two branches that lie approximately opposite in state space, each corresponding to one of the two spatial configurations of the juices. Within each branch, trajectories are further organized by choice: the split offer lies near the center, and trajectories for the two possible choices occupy opposite flanks while sharing the same spatial configuration.

**Figure 6.**
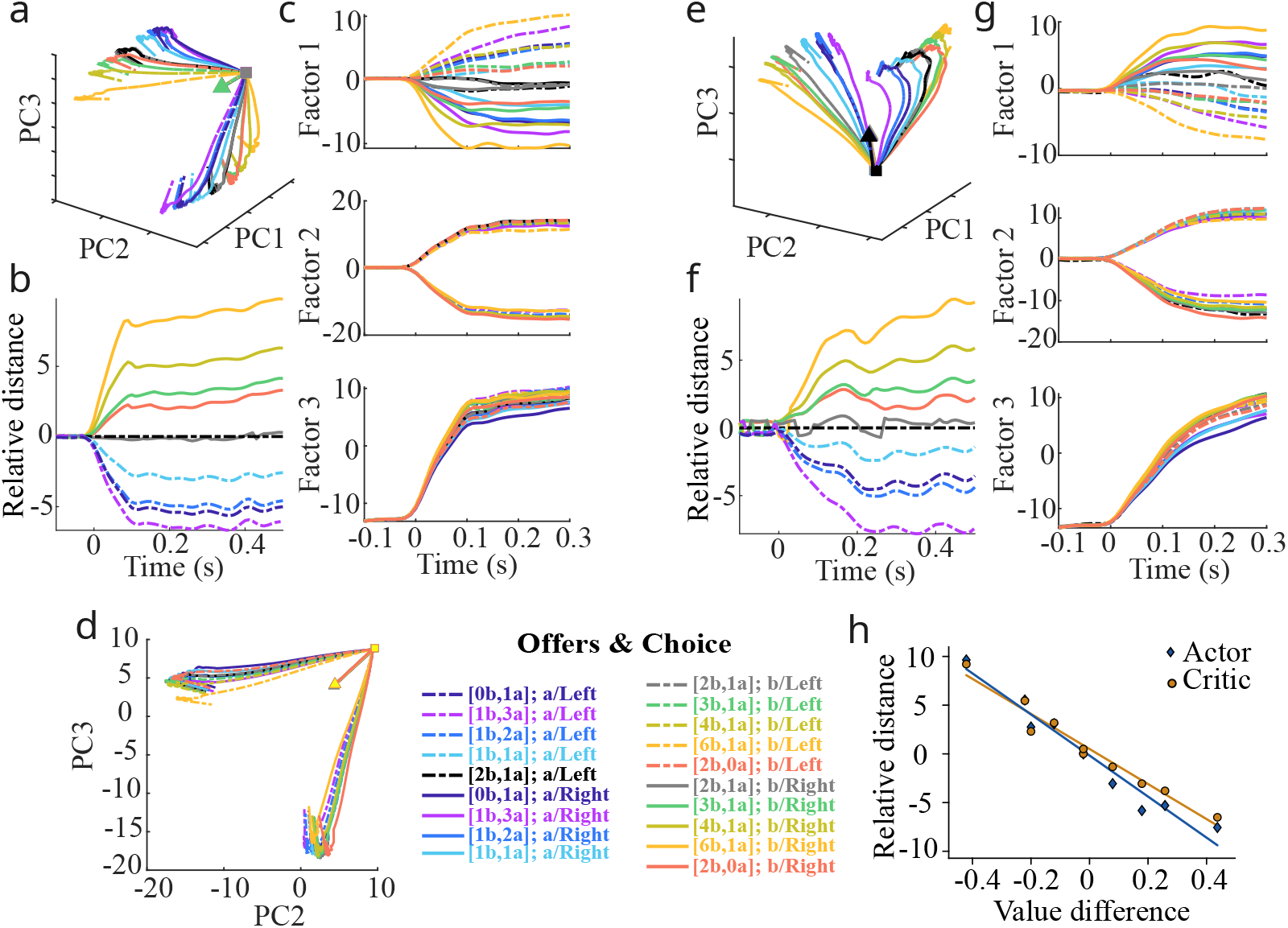
Dynamical organization of actor and critic networks. **(a)** Population neural activity is projected onto the first three PCs (86.68% of the total variance). Colors indicate offers; solid/dashed lines indicate Right/Left choices. Triangles mark task onset; squares mark offer onset (alignment point). Trajectories up to 1000 ms after offer onset form two branches corresponding to spatial configurations. **(b)** Relative distances among trajectories within the lower branch in (a) as a function of time, aligned to offer onset (*t* = 0). **(c)** Time courses of the first three PC factors. After rotating on the PC2-PC3 plane (Methods), factor 1 encodes value difference, factor 2 juice position, and factor 3 temporal evolution. **(d)** Neural state space defined by PC2 and PC3 allows visualization of the two branches corresponding to the two spatial arrangements. **(e–g)** Same analyses were performed for the critic network, yielding a 3-PC state space aligned to offer onset, which explained 86.89% of the variance. **(h)** Linear regression between the time-averaged relative distances and the value difference of each juice pair (blue: actor; brown: critic). Both networks show strong correlations 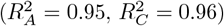.

This branching defines two spatial submanifolds that act as reference frames for subsequent computations. Within each submanifold, trajectories evolve along a value-related axis (approximately PC1). To determine which value variable best explains this organization, we computed the temporal mean of the relative distance between each trajectory and the split trajectory (200–1000 ms after offer onset) and regressed these distances against chosen action value, total action value, and action-value difference (Methods). The value difference between the two actions provided the best fit (*R*^2^ = 0.95; Fig. 6b,h, blue symbols; see also Supplementary Fig. 2). The interpretation of PC1 as encoding the action-value difference is further supported by the temporal evolution of the projection onto this axis (factor 1 in Fig. 6c, top): trajectories corresponding to opposite decisions for the split offer remain close, consistent with their location near the indifference point.

When projecting trajectories onto the PC2–PC3 plane (Fig. 6d), the line connecting the initial state to the offer onset appears as a symmetry axis. This symmetry reflects the interchangeability of left and right positions. Rotating this plane so that the symmetry axis aligns with PC3 (Supplementary Fig. 3) yields new components PC2^*′*^ and PC3^*′*^ with a clearer interpretation: the factor associated with PC2^*′*^ encodes spatial configuration, whereas the factor associated with PC3^*′*^ captures an approximately pure temporal signal, largely independent of offer identity and spatial configuration (Fig. 6c, middle and bottom). Thus, the three dominant principal components in the actor correspond to task-relevant dimensions: action-value difference (PC1), spatial position (PC2^*′*^), and temporal evolution (PC3^*′*^).

Although the critic was trained only to predict reward, its population activity exhibits a similar branching geometry after offer presentation (Fig. 6e). Again, trajectories split according to juice position, and within each branch they are organized along a value-related axis. Regressing the temporal mean of relative distances (Fig. 6f) against chosen juice value, total juice value, and juice-value difference revealed that the latter best explains the organization (*R*^2^ = 0.96; Fig. 6h; see also Supplementary Fig. 3). The projection onto PC1 illustrates this value structure (factor 1, Fig. 6g, top). Rotating the PC2–PC3 plane of the critic produces factors 2 and 3 with roles similar to those in the actor (Fig. 6g, middle and bottom), again linked to spatial configuration and time.

The similarity in trajectory structure between actor and critic can be understood mechanistically. The actor, trained to maximize reward through action selection, naturally develops an internal representation of action values organized along spatially defined branches. The critic receives this activity and is trained to predict reward; thus, it tends to encode juice-related value signals. Once the actor’s dynamics commit to one of the spatial branches, action values and juice values effectively coincide, so the signal transmitted to the critic carries structured value information that inherits the actor’s branching geometry. The critic’s population activity, in turn, reorients this inherited structure toward reward-predictive dimensions. Therefore, actor and critic share a common dynamical geometry imposed by input structure and task constraints, yet differ in computational function: one transforms values into actions, while the other evaluates outcomes in terms of predicted reward.

## Discussion

This work explores how interacting dorsal and ventral circuits may jointly implement valuation, confidence estimation, and action selection. Using a bimodular spiking actor–critic architecture trained on an economic choice task, we showed that the two modules develop distinct yet complementary representations that reflect their computational roles. The actor encodes action-value differences and spatial contingencies needed for committing to an action, whereas the critic represents object-related values predictive of future reward. The model reproduces behavioral performance and expresses several qualitative features reminiscent of neural responses observed across frontal and striatal regions.

Despite its simplicity, the model generates tuning types that parallel experimentally observed phenomena. Both modules contain neurons selective for object value, action value, total value, and value difference, consistent with signals reported in OFC, dlPFC, and dorsal striatum [22–24]. The critic also develops confidence-related signals whose dependence on the absolute value of the difference in the value of the two options resembles those described in neurophysiology [38]. Crucially, these signals emerge spontaneously from reinforcement learning rather than being imposed by architectural constraints or supervised objectives, supporting the view that object/action value, total value, choice, and confidence representations are distributed across fronto-striatal circuits.

It is noticeable that both modules exhibit predominant total-value signals. In primates, total value correlates with motivational variables such as vigor or reaction time [24, 34, 41, 42]. Although our model reliably produces total-value representations, it lacks these behavioral quantities that would allow us to determine whether these signals reflect motivational drive.

Low-dimensional analyses in state space revealed structured trajectories reflecting temporal evolution, spatial configuration, and value difference. Trajectories branched into two submanifolds corresponding to the spatial configuration; within each branch, activity evolved along axes related to decision formation. In the critic, this branching mirrored that of the actor because of unidirectional connectivity, explaining why critic activity remained position-dependent despite OFC position-invariance in primates.

A methodological contribution of this work is a novel procedure to map trained GRU networks onto spiking networks with gating mechanisms. Rather than embedding the spiking dynamics within the low-dimensional latent space of the original GRU system [29], our method constructs an expanded neural space in which the sRNN replicates the dynamics of the trained GRU. This expansion allows for a more distributed representation of latent variables across a larger spiking population, providing a framework in which task-relevant computations can be implemented using biologically inspired spiking dynamics.

We used a gated recurrent architecture, and there are compelling reasons to employ these models in computational neuroscience. Importantly, the original striatal gating theory was itself inspired by gated architectures [43], and converging evidence suggests that multiple gating-like mechanisms operate in the prefrontal cortex, functionally resembling those in artificial networks [27]. Gated architectures have also demonstrated strong empirical utility in modeling cognitive functions. For instance, J.-X. Wang et al. [19] showed that LSTM networks can learn meta-reinforcement learning strategies and replicate behavioral phenomena observed in animal studies. Similarly, Song et al. [17] used GRUs to model decision-making tasks and found results that aligned well with neural data, despite the abstract formulation of gating mechanisms. These examples suggest that gating plays a valuable computational role. Gating was also used by other authors [44]. Additionally, more biologically grounded alternatives to standard gating procedures have been proposed [45], offering directions for future computational models.

Our analyses reveal several limitations inherent to the simplified two-module architecture we investigated. Because valuation and choice formation are forced to coexist within a single ventral–dorsal division, the model cannot reproduce the diversity of empirical signatures observed in prefrontal cortex during value-based decision-making [22–24]. In particular, the current actor–critic system cannot develop an internal, categorical value-to-choice transformation such as the transition from graded object-value signals to an explicit, choice-selective representation, as reported in dorsolateral prefrontal cortex (dlPFC) in foraging tasks where values must be constructed from reward history [24]. Likewise, the model does not express a dominant population of chosen-object-value neurons in the critic module or chosen-action-value neurons in the actor module, unlike what is observed in OFC and dorsal striatum in explicit economic choice tasks [22, 23].

A related development concerns recent work showing that a single recurrent population trained end-to-end with reinforcement learning can reproduce several signatures of orbitofrontal value coding. In a relevant study, Battista et al. [21] trained a Dale-compliant vanilla RNN with PPO [46] across multiple economic-choice tasks and found that offer-value, chosen-value, and chosen-good signals emerged spontaneously within a single module. These results demonstrate that rich value representations can arise without imposing an explicit architectural separation between valuation and choice. However, because all computations—valuation, comparison, and choice—coexist within the same recurrent circuit, such models cannot exhibit a distinct and categorical value-to-choice transformation. Thus, while single-module RL approaches capture aspects of OFC physiology, they cannot address the missing computational step between graded object-value encoding and discrete choice formation observed in prefrontal cortex.

These discrepancies reflect the restricted computational scope that emerges when only two broad recurrent modules are trained jointly. A natural extension is to enrich the architecture by adding an intermediate recurrent module between a value-encoding network and an action-selecting network. When the three modules are trained end-to-end with a reinforcement learning algorithm in, e.g., the foraging task in [24], this intermediate module is expected to acquire the functional role of an *object-policy* network. Concretely, because reward depends on selecting the better object, policy-gradient optimization places pressure on this module to transform graded value inputs from the upstream critic-like network into a categorical object-choice signal. Through policy-gradient methods, the network could learn to implement a nonlinear competition among object-related representations, giving rise to choice-predictive and chosen-value signals analogous to those observed in dlPFC [24]. In parallel, the upstream value module can still develop mixed offer-value and chosen-value selectivity, consistent with OFC physiology in explicit choice tasks [22], while the downstream actor module learns to map the selected object to the appropriate spatial action that obtains it.

In this way, a trained three-module architecture provides a principled and flexible solution to the missing computational stage between valuation and choice. It allows the value-to-choice transformation to emerge naturally within ventral circuits prior to action selection, and can reproduce the distribution of value, chosen-value, and choice signals observed across OFC, dlPFC, and dorsal premotor/striatal areas, depending on training conditions and task demands.

Altogether, the present work shows that a minimal dorsal–ventral architecture can reproduce several hallmark features of value-based decision-making, spanning both object-related and action-related computations, while also revealing the necessity of intermediate processes, particularly value-to-choice transformations, in shaping emergent neural codes. Future extensions incorporating an explicit object-policy population, or more generally a multi-stage decision hierarchy, may help bridge the gap between model computations and the complex transformations observed in prefrontal and striatal circuits.

Crucially, we studied a framework that allows for scaling to such multi-modular spiking systems in a computationally tractable manner, enabling richer and more biologically grounded architectures.

## Acknowledgements

N.P., C.A., and L.S.-F. acknowledge that their work was conducted without external funding. J.F.-R. was funded by the European Union’s Horizon 2020 research and innovation programme (No. 857533), from the International Research Agendas Programme of the Foundation for Polish Science (No. MAB PLUS/2019/13), and from the Minister of Science and Higher Education “Support for the activity of Centers of Excellence established in Poland under Horizon 2020” (No. MEiN/2023/DIR/3796). The Polish high-performance computing infrastructure PLGrid (HPC Center: ACK Cyfronet AGH) was used to train the rate GRU networks (No. PLG/2025/018289).

## Author contributions

J.F.-R. and N.P. designed research; C.A. and J.F.-R. modified the code to train the rate networks; C.A. wrote the code to train and analyze the spiking networks with the help of J.F.-R.; C.A., J.F.-R., and N.P. analyzed the spiking networks with the help of L.S.-F.; L.S.-F. performed the state-space computations; C.A., L.S.-F., and N.P. analyzed the state-space results with the help of J.F.-R.; N.P. supervised research; C.A. and N.P. wrote the initial draft of the manuscript; C.A., J.F.-R., L.S.-F., and N.P. reviewed and edited the final version of the manuscript.

## Code and Data Availability

Matlab codes, as well as the raw data necessary for full replication of the figures, will be available upon publication.

## Methods

### Task description

The network was trained to perform a simplified adaptation of the economic choice task originally developed by Padoa-Schioppa and Assad [22], here implemented as a sequence of discrete trial phases. Each trial began with a fixation period of 750 ms, during which the agent was required to maintain fixation. Following this, one of nine predefined offer types (Table 1), specifying discrete amounts of juice a (*m*_*a*_) and juice b (*m*_*b*_), was presented for a variable duration between 1 and 2 seconds. During the offer presentation, the fixation cue remained on the screen. Afterward, a 750 ms decision period allowed the agent to choose between the two available juices.

At each time step of a trial, the agent made one of three possible actions: *fixate, left*, or *right*. During the offer period (Fig. 1d), selecting *left* or *right* prematurely (i.e., before the offer period ended) resulted in a constant negative reward *ρ* = −1 and immediate termination of the trial. During the decision period, choosing *left* or *right* yielded the amount of juice associated with the selected spatial location (either juice *a* or *b*). Juice amounts and their spatial locations were assigned randomly across trials from a predefined balanced set, ensuring a uniform distribution and preventing systematic spatial or value biases. If the policy network failed to choose either action before the end of the decision period, it again received a constant negative reward *ρ* = −1 and the trial was aborted.

The network’s output indicated a binary decision (choose *left* or *right*), and reward delivery was contingent on a probabilistic function of the simulated agent’s choice behavior. Over many trials, the agents’ preferences could be quantified using a psychometric function plotting the probability of choosing juice a over b as a function of the offers (Fig. 1e). The indifference point, defined as the ratio at which the model chooses either option with equal probability [17, 22], was computed by fitting the choice pattern with a sigmoid as a function of the normalized difference in offer magnitudes:

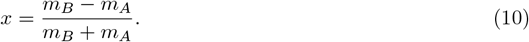

The sigmoid’s midpoint *x*^*∗*^, where the choice probability is 50%, defines the point of subjective equivalence. This value was then transformed into an indifference ratio:

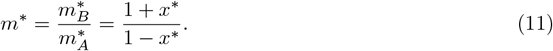

The indifference point, where both options are chosen with equal probability, provides an estimate of the relative value assigned to each juice.

### Rate policy and value networks with gated recurrent units

This section describes the method used to train the GRU networks before transferring their learned dynamics to the expanded gLIF networks. The procedure follows [17] in its entirety and is included here for the reader’s convenience. We defined the policy and value networks as gated recurrent unit (GRU) networks governed by the equations

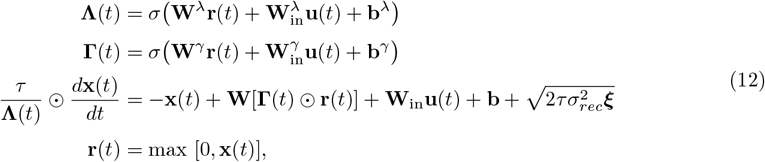

where the sigmoid function *σ*(*a*) = 1*/*(1 + *e*^−*a*^) operates the gates **Λ** and **Γ**, ⊙ is an element-wise multiplication, **b, b**^*λ*^, **b**^*γ*^ are biases, ***ξ*** are *N* Gaussian with zero mean and unit of variance noise processes where 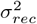 modulates the size of the noise, and *τ* = 100 ms is a time decay constant. Here, **W**^*λ*^, **W**^*γ*^, and **W** are the recurrent weight matrices of the hidden layer of the GRU network, and 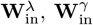, and **W**_in_ are the weight matrices of the input connections. **x**(*t*) is the current of the *M* neurons, and **r**(*t*) are the rectified-linear neural rates. The policy network received an *N ×* 7 matrix of inputs **u**^*π*^(*t*) that contained the task structure and dynamics (Fig. 1d). The value network received an *N ×* (*N* + 3) matrix of inputs **u**^*v*^(*t*) that contained a one-hot encoding of the action chosen by the policy network together with the rates **r**^*π*^(*t*) of its *N* units.

Denoting **Θ** as the set of plastic parameters of the policy network, the probability of selecting an action *a* at a given time was given by the softmax probability distribution

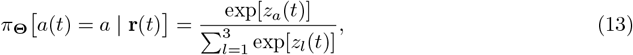

where

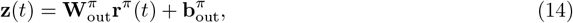

are the preferences of the possible actions, 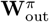 is a 3 *× M* matrix of output weights, and **b**_out_ is avector of output biases. Actions were sampled using inverse transform sampling.

Similarly, denoting **Φ** as the set of plastic parameters of the value network, its output is a scalar predicting the future return,

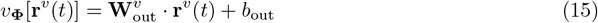

where 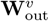 is an *M ×* 1 matrix of output weights and bias *b*_out_.

The policy network was trained to maximize the expected return [4],

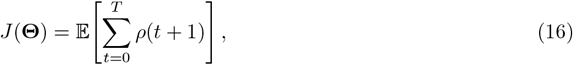

where *T* is the maximum trial duration. In practice, we optimized the loss

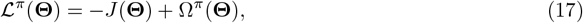

taking gradient steps along ∇_**Θ**_ ℒ^*π*^ (**Θ**) using the ADAM optimizer [47]. In a given trial, the gradient of the expected return was computed using the REINFORCE estimator [25]

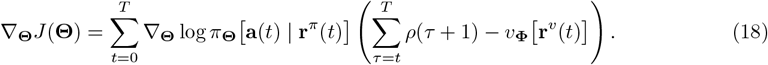

The eligibility term ∇_**Θ**_ log *π*_**Θ**_(**a**_*t*_ | **u**_1:*t*_) and the entropy-regularization gradient ∇_**Θ**_Ω^*π*^(**Θ**) were computed using backpropagation through time [48].

The value network was trained to minimize the mean-squared return prediction error [4],

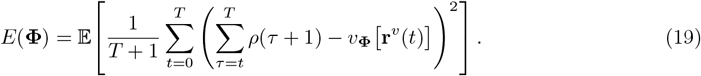

More precisely, we optimized the loss

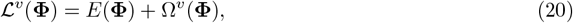

taking steps along ∇_**F**_ ℒ ^*v*^ (**F**) using the ADAM optimizer [47]. Both the prediction-error gradient ∇_**F**_*E*(**F**) and the entropy-regularization gradient ∇_**F**_Ω^*v*^(**F**) were obtained using backpropagation through time [48].

The training was conducted in Python 2.7.18 using the code provided by Song et al. [17], with identical parameters. Each network consisted of 100 gated recurrent units (GRUs) and was trained with a temporal resolution of *dt* = 10 ms. Gradient updates were performed after all the offers in Table 1 had been presented. All weights were initialized following the same procedures as in Song et al. [17]. Recurrent weights were further rescaled to achieve a spectral radius *R* = 2, defined as the modulus of the largest eigenvalue [49]. This value placed the dynamics in the chaotic regime (*R >* 1) but close to the stability boundary, thereby enhancing variability and exploration during learning [50]. Only the recurrent and output weights for both the actor and critic networks underwent plastic changes along the training process, i.e., 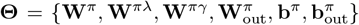 and 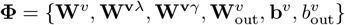.

Importantly, the training termination criterion was modified from fixed performance thresholds [17] to a convergence tolerance between the agent’s decision and choice correctness probabilities. For every consecutive batch of *N*_*validation*_ validation trials, the probability of choosing the highest valued offer *p* was estimated using an exponential running average of the history of choices,

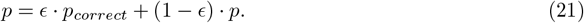

The probability of a choice being correct was estimated within each training batch by computing the fraction of trials that resulted in the choice with the highest amount of juice *p*_*correct*_ = *N*_*correct*_*/N*_*validation*_. The training ended once the behavior of the network in validation trials remained stable around an arbitrary value |*p* − *p*_*correct*_| ≤ *δ*, determined solely by the network choices, ultimately yielding an average reward rate. The exponential decay rate was set to *ϵ* = 0.2 and the tolerance to *δ* = 0.01. We initially set *p* to −1.

### An actor-critic system of gated leaky integrate-and-fire neurons

We implemented a spiking system of two interconnected modules mirroring the policy and value GRUs in Eq. 12. Each module contained *N* gated leaky integrate-and-fire (gLIF) units whose subthreshold dynamics were

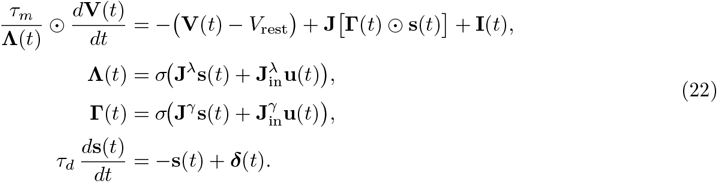

Here, the *N × N* weights **J, J**^*λ*^, **J**^*γ*^ and the *N × N*_in_ input weights **J**^*λ*^, **J**^*γ*^ are given by Eqs. 6 and 7. Synaptic currents **s**(*t*) were modeled with a single-exponential filter of presynaptic spikes, with ***δ***(*t*) denoting an *N ×* 1 vector of spike impulses of components 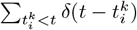, where 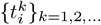 are the presynaptic spiking times of neuron *i*. Each gLIF unit emitted a spike when *V*_*i*_(*t*) ≥ *V*_*th*_. Immediately after spiking, *V*_*i*_(*t*) was reset to *V*_*rest*_ and held for a 10 ms refractory period (one simulation step). We used *τ*_*m*_ = 10 ms for the membrane constant, *τ*_*d*_ = *τ* = 100 ms for the synaptic decay time, *V*_*th*_ = −55 mV for the spiking threshold, and *V*_*rest*_ = −65 mV for the resting potential.

The input current to each neuron comprised three components

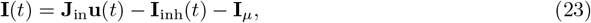

where **J**_in_ is a matrix of *N × N* input weights given by Eq. 7, and **u**(*t*) encoded the task structure and dynamics. To regulate overall network activity [28, 29], we introduced a global inhibition current

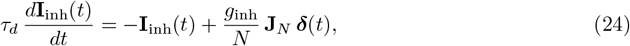

where **J**_*N*_ is the *N × N* all-ones matrix and *g*_inh_ = 0.28. This current rose with global network spiking and was subtracted from the input, acting as a stabilizing mechanism that prevented runaway excitation and network-wide synchrony.

Lastly, the mean input current **I**_*μ*_ to each neuron was defined as the average intensity across time and trials in the absence of input; i.e., **u**(*t*) = 0. More precisely, we simulated 1500 trials, obtained the total intensity — including the recurrent synaptic currents and the global inhibition, and computed its average value across time and trials. This procedure eliminated the need to consider the constant biases present in the dynamics of the GRU networks, since their contribution would be canceled out during the averaging. Subtracting **I**_*μ*_ centered the membrane potential near *V*_*rest*_, promoting an irregular firing regime. Inputs did not model the fixation cue nor the *fixate* action (Fig. 1c, d).

### Network Size Expansion

The expansion from *M* to *N > M* units not only increases the representational capacity of the system but also preserves, up to a time scaling, the original dynamics of the GRU networks within a subspace of the higher-dimensional gLIF systems. This preservation is a direct consequence of the use of an orthonormal expansion matrix **U** satisfying **U**^⊤^**U** = **I**_*M*_. Specifically, since the dynamics of the gLIF networks are constructed in the image of the GRU activity under the transformation **U**, the evolution of the population activity within this subspace remains unchanged, up to a temporal scaling and noise-related perturbations (Supplementary Text S1.1). The gLIF dynamics thus replicate the original GRU trajectories in the embedded subspace, ensuring a faithful transfer of function.

### Simulation of gLIF networks

Simulations were performed in MATLAB (MathWorks) using gLIF networks composed of *N* = 400 neurons, obtained by expanding the original RNNs with *M* = 100 neurons. Networks were simulated over 1,500 trials. Similarly, the bias term was suppressed, as its effect was already accounted for by the mean current with opposite sign. Continuous RNNs outputs over time were transformed into discrete trial-level measures for both networks.

### Learning transfer from GRU to gLIF networks

Learning transfer from the continuous GRU networks to the corresponding gLIF spiking networks was performed using the actor and critic loss functions (Eqs. 8 and 9, respectively).

For the actor network, trial-by-trial readouts were constructed from the gLIF activity during the offer-period window (350 ms after offer onset, spanning 700 ms) by fitting the actor GRU outputs with MATLAB’s fitlm function. Cross-validation was used to estimate regression weights, and the sign of the resulting predicted preference determined the selected action for each trial.

For the critic network, readouts were trained on GRU-predicted rewards at the time of choice. Firing rates from the offer-period window were fitted using `fitlm` with cross-validation, and the resulting weights were used to compute trial-wise predicted rewards.

These readouts were used to determine approximate ranges of the scaling parameters *κ*_*η*_ (*η* = *A, C*) where learning transfer was satisfactory. Fine-tuning was then performed using, for the current value of *κ*_*η*_, the *diff* populations of each network, which are selectively tuned to action- and object-value differences. Population activity defined a decision variable (see below), from which behavioral choices were inferred and used to fine-tune the scaling parameters via the actor loss in Eq. 8.

### Neural activity

Firing rates were computed using a causal boxcar filter with 150 ms windows and 10 ms steps. The offer-period analysis window started 350 ms after offer onset and spanned 700 ms to capture peak activity. A pre-stimulus window of 200 ms was defined during the noisy baseline period, finishing 350 ms before offer onset.

z-score normalization was applied neuron-by-neuron across trials, separately for pre-tuning and post-tuning activity. Normalization was computed within each condition, defined by the combination of offer and choice, to standardize firing rates and control for baseline variability, thereby facilitating comparison across neurons.

Raster plots were generated from z-scored firing rates and spike trains across all offer types. For each offer condition, 50 random trials were selected and grouped by value-based variables (value difference, offer value a/b, or offer sum).

### Actor network regression in terms of action values

For each actor neuron, the firing rate in the offer-period window *S*_*A*_ was regressed onto choice and action value predictors: the chosen action 𝒜 (+1 for right, −1 for left), action values 𝒜 𝒱_*R*_ and 𝒜 𝒱_*L*_, defined as the reward magnitudes associated with the juices offered at the right and left [23], and the juice position regressor 𝒥_*pos*_ (1 if juice *a* on the left, 0 otherwise). The model was specified as

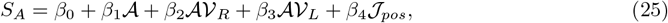

where *β*_*i*_ denote regression coefficients. Neurons were classified as encoding action values if the overall model was statistically significant (F-test, *p <* 0.01).

### Critic network regression in terms of juice values

For each critic neuron, the firing rate in the offer-period window *S*_*C*_ was regressed onto juice (object) values, 𝒱_*a*_ and 𝒱_*b*_, as well as the juice position regressor 𝒥_*pos*_. These values were defined as 𝒱_*a*_ = 0.1 *m*_*a*_ *m*^*∗*^ and 𝒱_*b*_ = 0.1 *m*_*b*_, where *m*_*a*_ and *m*_*b*_ denote juice quantities, and *m*^*∗*^ is the indifference point estimated from the RNNs model. The regression model was specified as

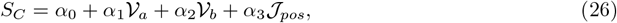

where *α*_*i*_ are the fitted regression coefficients. Neurons were classified as encoding juice values if the overall model was statistically significant (F-test, *p <* 0.01).

### Compound partial responses

To isolate the contributions of regressors relevant for decision computation, we computed compound partial responses for each neuron *m*. In the actor network, this includes only the action value regressors (𝒜 𝒱_*R*_, 𝒜 𝒱_*L*_)

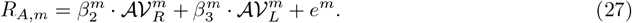

In the critic network, compound partial responses include the object value regressors (𝒱_*a*_, 𝒱_*b*_)

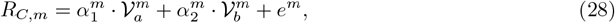

where *e*^*m*^ denotes residual variance not explained by the selected regressors.

### Decision variables from neural populations

Decision variables (DV) were constructed from population-averaged compound partial responses. In each network, neurons for which the relevant pair of regression coefficients (action values in the actor, juice values in the critic) were both significant with opposite signs were grouped into two populations corresponding to *diff* population. Neurons with a positive first coefficient (*β*_2_ or *α*_1_) and a negative second coefficient (*β*_3_ or *α*_2_) comprised the *plus* population (Sector 7), whereas those with the opposite signs formed the *minus* population (Sector 3). Population-averaged compound partial responses were computed as

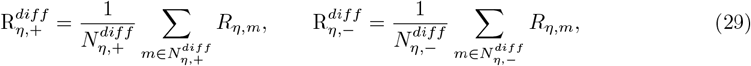

where *η* = *A, C* indicates the actor or critic network. While either population could be used independently to make a choice, the optimal readout is expected to be given by the difference [51]

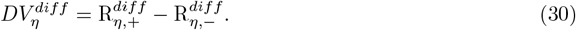

We further extended the analysis to neural populations in which only one regressor was significant. *Plus* and *minus* populations were defined by the sign of the significant coefficient (positive: *plus*, negative: *minus*). For each population, the decision variable was computed as the difference between *plus* and *minus* compound partial responses. In the actor network, these population DVs were computed for *AV R* and *AV L*, and in the critic network for *OV a* and *OV b*. The final decision variables capturing action value and object value contrasts were then obtained as the difference between the respective population DVs:

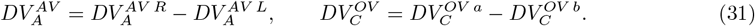

For the actor network, *DV*_*A*_ *>* 0 predicts a right action, while *DV*_*A*_ *<* 0 predicts a left action. For the critic network, *DV*_*C*_ *>* 0 predicts choice of juice *a*, while *DV*_*C*_ *<* 0 predicts choice of juice *b*.

### Population analysis of action value sensitivity

To quantify the influence of right and left action values on neuronal activity, we computed partial residuals from the actor regression model. For each neuron *m* and regressor *i*, the partial residual 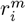 was defined as

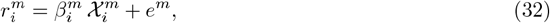

where 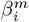 is the corresponding regression coefficient, 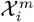 is the regressor value, and *e*^*m*^ denotes the residual variance unexplained by the selected regressor.

To assess sensitivity to action value, for each trial we identified the regressor corresponding to the larger action value (𝒜 𝒱_*R*_, 𝒜 𝒱_*L*_) and averaged the associated partial responses across trials, separately for each chosen action (*R* or *L*). Residuals from *minus* populations were averaged with an inverted sign relative to their corresponding *plus* populations.

### Discriminating between Total Value and Chosen Value Coding

To determine whether neurons in the *sum* sectors of the actor (critic) encoded total action value or chosen action value (total juice value or chosen juice value), we applied the procedure described in [22]. For each neuron, we performed two separate linear regressions of the firing rates on total action value and chosen action value (total juice value or chosen juice value), respectively, using data from the offer-period window. The coefficients of determination (*R*^2^) obtained from both fits were then compared. When both regressions yielded significant slopes, *R*^2^ values from the two regressions were compared using a Wilcoxon signed-rank test to assess which variable better explained the neuron’s activity. This analysis allowed us to classify neurons as encoding either total or chosen action value (total juice value or chosen juice value), and to determine which of these representations predominated in the population.

More precisely, in the actor network linear regressions were performed using the total action value (𝒜 𝒱_*a*_ + 𝒜 𝒱_*b*_) or the chosen action value 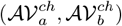 as predictors. In the critic network, we followed a similar procedure, performing separate linear regressions using either the total value, sum of both juice options (𝒱_*a*_ + 𝒱_*b*_), or the chosen juice value, 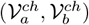, as predictors. Neural activity was z-scored across trials within each offer–choice condition. Regressions were then computed at the single-neuron level and at the population level using mean activity, with MATLAB’s `fitlm` function. For the actor network, this analysis was performed separately for each spatial configuration.

### Bayesian logistic regression for trial-wise confidence

To quantify decision confidence from population activity, we used a Bayesian framework to estimate trial-wise statistical confidence. For each trial *i*, let **s**_*i*_ denote a 1 *×* (*N* + 1) vector containing the spike rates of the *N* neurons plus a constant term, and let *y*_*i*_ ∈ *{*0, 1*}* indicate whether the agent chose the option with the higher reward. Given the training data 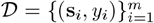, the optimal predictive probability for a new observation **s** of *N* neural spike rates is

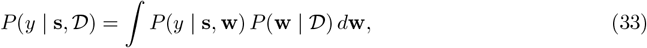

where **w** is an (*N* + 1) *×* 1 weight vector including a bias. We defined the Bayesian decision confidence [36] as the probability of making a correct choice, *P* (*y* = 1 | **s**, 𝒟).

We adopted a logistic function for the generative choice model because it naturally fitted the binary nature of the economic choice task.

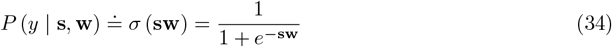

To compute the posterior distribution of the weights, we assumed a Gaussian prior

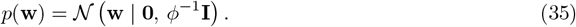

which, using the Laplace approximation, allowed us to write the posterior distribution also as a Gaussian [40, chapter 4, p. 218],

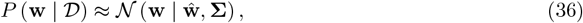

centered around the maximum a posteriori (MAP) estimate of the log-posterior,

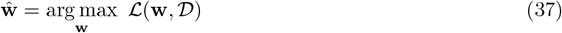

and with covariance matrix

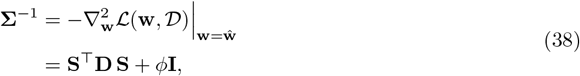

where

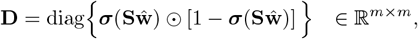

and **S** is n *m ×* (*N* + 1) a matrix containing the spiking rates of the neurons and constant for all *m* trials. Supposing i.i.d. samples, the likelihood of the data is

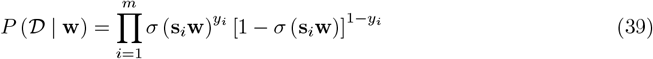

and

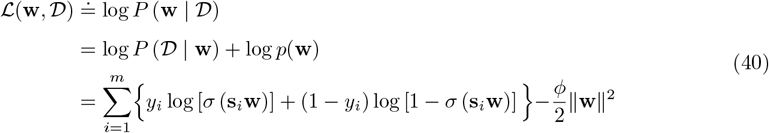

is the log-posterior of the weights given the training data 𝒟.

Because of the sigmoid function, there is no closed form for the MAP estimate in Eq. 37. Consequently, we employed the Newton-Raphson method to minimize the negative log-posterior and obtain a numerical solution iteratively. For that, we set the initial weights **ŵ** (i.e., prior mean in Eq. 35), calculated the corresponding covariance matrix **Σ** using Eq. 38, and updated the MAP estimate according to

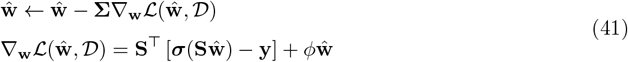

where **y** is an *m ×* 1 column vector containing the outcomes for all trials. Then, the covariance matrix **S** and gradient ∇_**w**_ℒ were recomputed with the updated values of **ŵ**, and the procedure was repeated until ||**S**∇_**w**_ℒ (**ŵ**, *D*)|| *<* 1*e*^−6^.

Finally, the decision confidence Eq. 33 can be calculated by substituting the two distributions in Eqs. 34 and 36,

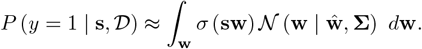

This integral does not admit a closed-form solution, but it can be accurately approximated to yield an analytic expression for the Bayesian decision confidence [40]

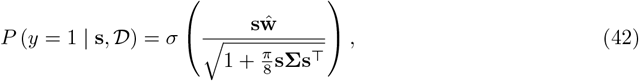

where **ŵ** and **S** are the solutions from the Newton-Raphson procedure. The precision parameter *ϕ* ∈ [1, 200] in Eq. 35 was tuned by selecting the value that maximized the correlation between Bayesian decision confidence and the absolute offer-value difference |Δ*V* |. The results were robust to the resulting choice of *ϕ* = 200; varying this parameter did not affect convergence or the inferred confidence values.

### Principal Component Analysis

To uncover low-dimensional neural state spaces, we performed principal component analysis (PCA) on trial-averaged population firing rates conditioned on offer type: juice pair presented and position of both juices. For the quantitative assessment of trajectory geometry (Fig. 6), we constructed neural state spaces using the first three principal components, which retained the 87% of the total variance in each network.

For each principal component, we defined a corresponding factor as the projection of the z-scored neuronal activity onto that component (Fig. 6c and g). Factors were computed separately for each condition and time point. To enhance interpretability, the PC2–PC3 plane was rotated by a fixed angle (see Supplementary Fig. 3).

### Relative distances between state-space trajectories

To quantify the separation between neural trajectories, we applied the Kinematic analysis of Neural Trajectories (KiNeT) method [52]. This approach compares a given trajectory (*j*) to a selected reference trajectory (*ref*) by evaluating their relative distances over time. At each time point *t*, we identified the neural state of the reference trajectory, *S*_ref_(*t*), and searched for the closest state on trajectory *j*, denoted *S*_*j*_(*t*), in terms of Euclidean distance. The relative distance at time *t* was then defined as

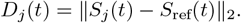

All distance calculations were performed within the PC spaces described above. Reference trajectories were the split trajectories: for trajectories ending in choice *a*, the reference was the corresponding split trajectory with choice *a*, and analogously for trajectories with choice *b*. Thus, *j* = 1, …, *N*_*a*_ or *j* = 1, …, *N*_*b*_, where *N*_*a*_ and *N*_*b*_ denote the number of trajectories with choice *a* or *b*, respectively. To characterize the geometry of the trajectories, we computed the temporal mean, *D*_*j*_, of the relative distances during a temporal window extending from 200 to 1000 ms after offer onset (Fig. 6b,f).

## S1 Supplementary Texts

### S1.1 Dynamics of the Expanded Spiking Network

Here, we present a more formal discussion of the dynamics of the expanded spiking network. For simplicity, we ignore the gating variables and consider a linearized rate model. Let **r**(*t*) denote the *M*-component vector of activities in the original rate network, governed by

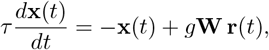

where *τ* is the characteristic time constant and **W** is the *M × M* recurrent coupling matrix. The gain factor *g* simply rescales the eigenvalues of **W** and can therefore be absorbed into the coupling matrix without loss of generality, i.e., *g***W** → **W**.

To construct the spiking counterpart, we define an expanded network of *N* neurons with membrane potentials collected in the *N*-component vector **V**(*t*). Their subthreshold dynamics follow

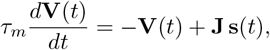

where *τ*_*m*_ is the membrane time constant, **J** is the *N × N* synaptic weight matrix, and **s**(*t*) is the vector of synaptically filtered spike trains. Each component *s*_*i*_(*t*), *i* = 1, …, *N*, is obtained from the sequence of presynaptic spiking times 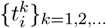 through exponential filtering,

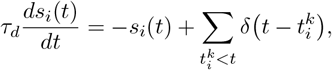

with *τ*_*d*_ the synaptic decay constant. The connectivity matrix **J** is defined as

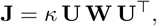

where **U** is an *N × M* orthonormal matrix satisfying **U**^⊤^**U** = **I**_*M*_, and *κ* is a scaling factor that will control the overall firing rate level of the spiking network.

The low-dimensional activity of the rate model is embedded into the spiking network via

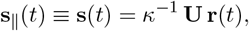

so that **s**∥(*t*) lies within the subspace spanned by the columns of **U**. Any spiking activity vector **S**(*t*) can be decomposed as

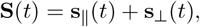

where **s**_⊥_(*t*) is orthogonal to the columns of **U**. Within the embedded subspace, the membrane potential dynamics satisfy

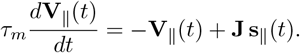

Using **s**∥(*t*) = *κ*^−1^**Ur**(*t*) and **J** = *κ***UWU**^⊤^, we obtain

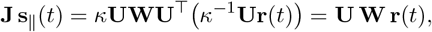

so that the factor *κ* cancels exactly in the recurrent term. Projecting onto the embedded subspace by multiplying on the left by **U**^⊤^ and defining **x**(*t*) = **U**^⊤^**V**∥(*t*), we obtain

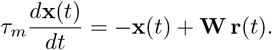

The eigenvalues governing the effective dynamics in this subspace are the *M* eigenvalues *λ*_*i*_ of **W**; the scaling factor *κ* drops out of the effective operator and thus does not affect the stability of the embedded latent dynamics. Instead, *κ* controls the overall magnitude of the spiking activity **s** associated with a given latent trajectory **r**(*t*).

For the orthogonal component **s**_⊥_(*t*), one obtains

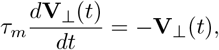

because **U**^⊤^**s**_⊥_(*t*) = **0** implies **J s**_⊥_(*t*) = **0**; that is, the remaining *N* − *M* eigenvalues of **J** are zero. This results in an exponential decay toward zero. These components do not contribute to the recurrent feedback and remain dynamically silent, except for fluctuations induced by noise or spiking variability.

### S1.2 Decision Confidence as a function of |Δ*V* |

We define confidence as the probability *P*_0_ that the agent selects the option with the higher subjective value. This quantity can be related to |Δ*V* |, the absolute difference in subjective value between the two options, using standard arguments from Signal Detection Theory. Let *V*_1_ and *V*_2_ denote the true values of two offers, and assume that the agent makes noisy estimates

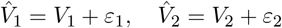

where *ε*_1_, *ε*_2_ ∼ 𝒩 (0, *σ*^2^) are independent noise terms with variance *σ*. The agent chooses the option with the higher estimated value. Define the decision variable

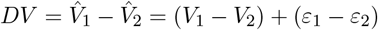

so that *DV* ∼ 𝒩 (Δ*V*, 2*σ*^2^), with Δ*V* = *V*_1_ − *V*_2_. The probability of choosing the better offer (i.e., the one with the higher true value) is

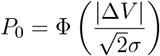

where Φ(*·*) denotes the standard normal cumulative distribution function.

## S2 Supplementary Figures

**Supplementary Figure 1.**
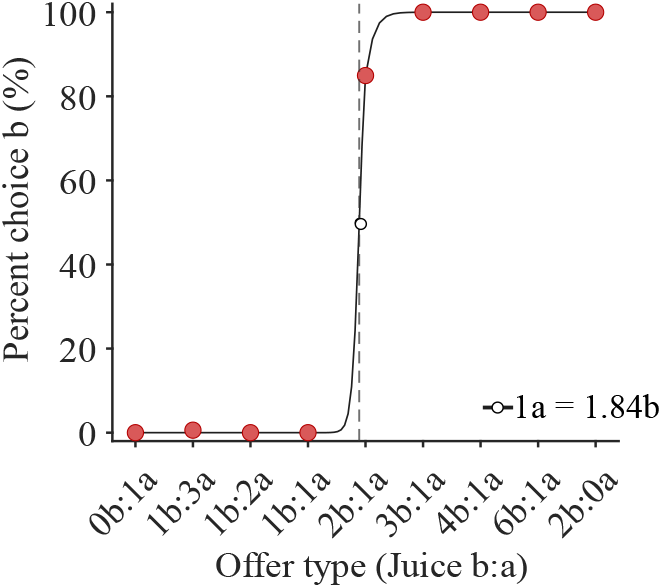
Choice pattern from the preference readout of the gLIF actor network. Percentage of trials where the gLIF actor linear readout preferred juice b. The indifference point was *m*^*∗*^ = 1.84 (solid black; loss *L*_*A*_ = −0.3276).

**Supplementary Figure 2.**
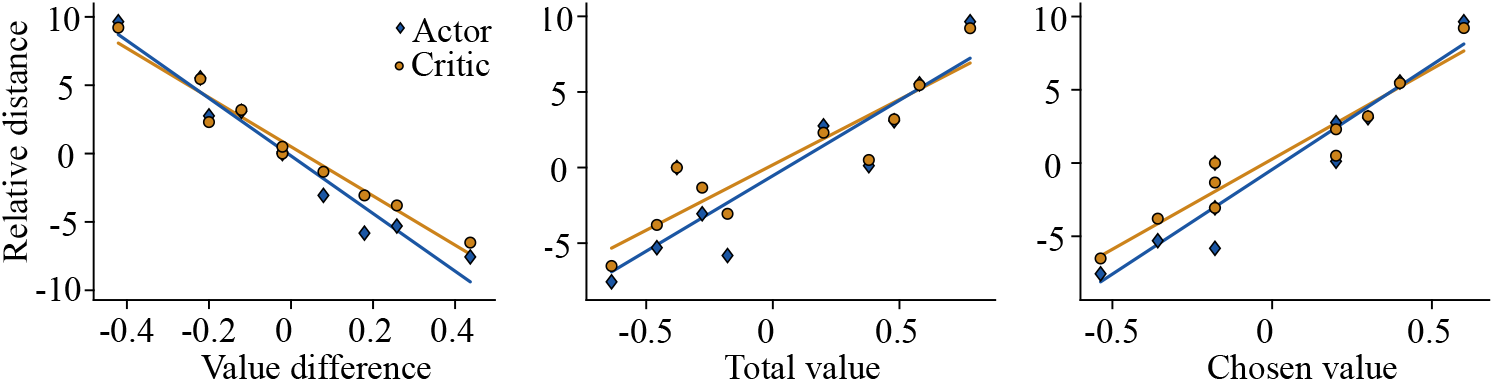
Relative distances between neural trajectories encode the value difference between juice quantities. Related to Fig. 6. For both actor (blue diamonds) and critic (orange circles) networks, linear regression of the temporal mean of the relative distances (relative to the split offer) as a function of the value difference (left), the total value (middle), and the chosen value (right). The goodness of fit is given by *R*^2^. (Left) Value difference: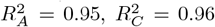. (Middle) Total value: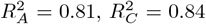. (Right) Chosen value: 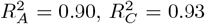. Among all encodings, the value difference better explains the variance in both networks. Note that, although the value-difference and chosen-value regressions have similar *R*^2^ values, in the chosen-value case three distances correspond to the same chosen value (which is not evident from the trajectories), whereas for value difference these distances are ordered along a sequence.

**Supplementary Figure 3.**
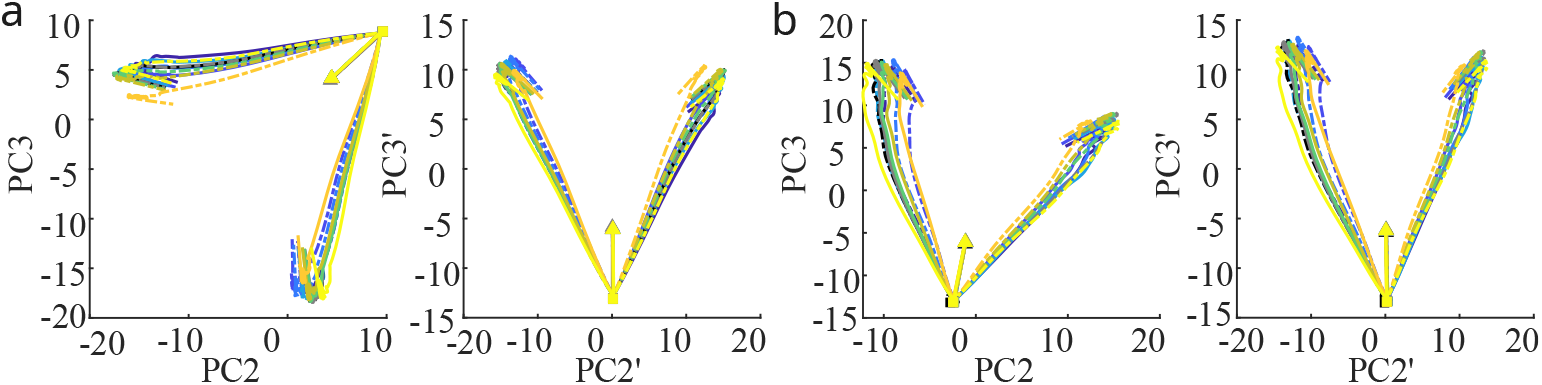
Geometry of PC2–PC3 projections before and after rotation. Related to Fig. 6. Projecting population trajectories onto the PC2–PC3 plane reveals an approximate symmetry between left and right offers, which can be captured by a single axis. We rotated this plane so that this axis aligns with PC3, defining new components PC2’ and PC3’ with a clearer task-related interpretation. (**a**) Actor network (left, original PC2–PC3 plane; right, rotated PC2’–PC3’ plane). (**b**) Critic network (left, original PC2–PC3 plane; right, rotated PC2’–PC3’ plane). In both networks, PC2’ primarily captures spatial configuration, whereas PC3’ reflects temporal evolution during the trial. Triangles mark task onset and squares indicate offer onset (alignment point). Trajectories are shown up to 1000 ms after offer onset. Color and line codes are the same as in Fig. 6.

## S3 Supplementary Tables

**Supplementary Table 1.** Distribution of value-related tuning in the critic gLIF network across spatial configurations. Neurons were first assigned to tuning sectors using all trials, irrespective of juice location (Fig. 4, Main Text). We then reassessed tuning using only trials in which juice *a* appeared consistently on the right or the left. The columns *Juice a Right* and *Juice a Left* report the number (and percentage) of neurons that preserved their original tuning under each fixed spatial configuration. The column *Juice a Right and Left* lists neurons that maintained the same tuning in both configurations independently, whereas *Juice a Right or Left* counts neurons that preserved tuning in at least one configuration (i.e., the union of the two single–location sets, excluding those counted in both). The fraction of neurons whose value tuning depended on spatial position (6th column) far exceeded the fraction that maintained the same tuning across both configurations (5th column).

| Tuning type | # Neurons (%) | # Neurons (%) | # Neurons (%) | # Neurons (%) | # Neurons (%) |
| --- | --- | --- | --- | --- | --- |
|  | *All trials* | *Juice a Right* | *Juice a Left* | *Juice a Right and Left* | *Juice a Right or Left* |
| *sum* (+) | 49 | 14 (28.6%) | 21 (42.9%) | 6 (12.2%) | 29 (59.3%) |
| *sum* (-) | 55 | 16 (29.1%) | 23 (41.8%) | 3 (5.5%) | 36 (65.4%) |
| *diff* (+) | 42 | 17 (40.5%) | 19 (45.2%) | 1 (2.4%) | 35 (83.3%) |
| *diff* (-) | 39 | 14 (35.9%) | 21 (53.9%) | 0 (0.0%) | 35 (89.8%) |
| *OV a* (+) | 38 | 21 (55.3%) | 9 (23.7%) | 4 (10.5%) | 26 (68.5%) |
| *OV a* (-) | 38 | 16 (42.1%) | 14 (36.8%) | 0 (0.0%) | 30 (78.9%) |
| *OV b* (+) | 64 | 41 (64.1%) | 21 (32.8%) | 3 (4.7%) | 59 (92.2%) |
| *OV b* (-) | 58 | 34 (58.6%) | 21 (36.2%) | 6 (10.3%) | 49 (84.5%) |

**Supplementary Table 2.** Distribution of action-value-related tuning in the actor gLIF network across spatial configurations. Neurons were first assigned to tuning sectors using all trials, irrespective of juice location (Fig. 3, Main Text). We then reassessed tuning using only trials in which juice *a* appeared consistently on the right or the left. The columns *Juice a Right* and *Juice a Left* report the number (and percentage) of neurons that preserved their original tuning under each fixed spatial configuration. The column *Juice a Right and Left* lists neurons that maintained the same tuning in both configurations independently, whereas *Juice a Right or Left* counts neurons that preserved tuning in at least one configuration (i.e., the union of the two single–location sets, excluding those counted in both). The fraction of neurons whose value tuning depended on spatial position (6th column) exceeded the fraction that maintained the same tuning across both configurations (5th column; albeit in slightly reduced amounts compared to the critic gLIF network.

| Tuning type | # Neurons (%) | # Neurons (%) | # Neurons (%) | # Neurons (%) | # Neurons (%) |
| --- | --- | --- | --- | --- | --- |
|  | *All trials* | *Juice a Right* | *Juice a Left* | *Juice a Right and Left* | *Juice a Right or Left* |
| *sum* (+) | 35 | 17 (48.6%) | 16 (45.7%) | 7 (20.0%) | 26 (74.3%) |
| *sum* (-) | 34 | 20 (58.8%) | 12 (35.3%) | 3 (8.8%) | 29 (85.3%) |
| *diff* (+) | 53 | 22 (41.5%) | 21 (39.6%) | 10 (18.9%) | 33 (62.3%) |
| *diff* (-) | 46 | 24 (52.2%) | 19 (41.3%) | 8 (17.4%) | 35 (76.1%) |
| *AV R* (+) | 57 | 20 (35.1%) | 42 (73.7%) | 15 (26.3%) | 47 (82.5%) |
| *AV R* (-) | 40 | 10 (25.0%) | 20 (50.0%) | 4 (10.0%) | 26 (65.0%) |
| *AV L* (+) | 65 | 54 (83.1%) | 19 (29.2%) | 17 (26.2%) | 56 (86.2%) |
| *AV L* (-) | 60 | 37 (61.7%) | 20 (33.3%) | 8 (13.3%) | 49 (81.7%) |

